# BrainRoute integrates machine learning prediction with community-driven verification of blood-brain barrier permeability

**DOI:** 10.64898/2026.11.19.767839

**Authors:** Soham A. Shirolkar, Lewis T. Bueh, Leah W. Cerere, Noura E. Ahmed, Georges Somé, Olaitan I. Awe

## Abstract

The blood-brain barrier (BBB) is a highly selective barrier that restricts the entry of many therapeutic agents into the brain. The lack of efficient methods for BBB permeability research and data management remains a significant challenge for central nervous system drug-development pipelines. Although BBB permeability data are available from multiple open sources, few platforms integrate these datasets with molecular exploration, predictive tools, and a structured process for community-supported data verification.

Here, we present BrainRoute, an open platform for BBB molecular data collection, exploration, prediction, and community-supported verification. BrainRoute provides searchable and filterable molecular records, physicochemical and structural information, source-linked data, single-molecule and batch prediction, and a structured workflow through which users can submit additional evidence for review. The prediction tool model-development cohort contained 7,888 unique molecules, including 5,424 BBB+ and 2,464 BBB-molecules. Six machine-learning classifiers were evaluated using PaDEL descriptors, Morgan fingerprints, frozen ChemBERTa embeddings, and combined feature representations. The three selected models were integrated into the platform as a probability-averaged ensemble. The primary five-fold scaffold-grouped analysis achieved a best mean balanced accuracy of 0.783 ± 0.026. On an independent external set containing 410 non-overlapping molecules, the deployed ensemble achieved a balanced accuracy of 0.659 (95% CI 0.600–0.712) and an AUPRC of 0.670 (95% CI 0.583–0.752). BrainRoute serves as a platform for exploring BBB-related molecular data and computational predictions while supporting the continued growth and verification of the underlying dataset. BrainRoute is freely accessible online (https://github.com/omicscodeathon/brainroutedb).

## 1. Introduction

The blood-brain barrier (BBB) is a highly selective and dynamic physiological interface that regulates the transport of molecules, ions, and cells between the systemic circulation and the central nervous system (CNS) (Kadry et al., 2020). It is primarily formed by specialized brain microvascular endothelial cells joined by tight junctions, which severely restrict paracellular diffusion and maintain CNS homeostasis. The BBB is supported by pericytes and astrocytes (Kadry et al., 2020). Pericytes are mural cells embedded within the vascular basement membrane that surround brain capillaries and are essential for BBB integrity, regulating endothelial tight junctions, vascular stability, angiogenesis, immune cell trafficking, and suppressing nonspecific vesicular transport (Armulik et al., 2010; Daneman et al., 2010). Astrocytes, a major glial cell type in the CNS, extend endfeet that ensheathe most of the brain microvasculature and modulate BBB permeability, ionic and neurotransmitter homeostasis, and long-term barrier maintenance through metabolic support and secreted signaling factors (Kadry et al., 2020). Furthermore, BBB function can also vary across toxicological exposures, neurodegenerative disease models, and brain tumors (Belykh et al., 2020; Katt et al., 2019; Molot et al., 2021; Zheng, 2001)

BBB permeability is a decisive factor in determining whether a therapeutic compound can reach its CNS target. At the same time, for many drugs, limited BBB penetration is also desirable, as avoiding entry into the brain can reduce unwanted central nervous system side effects; for example, second-generation antihistamines are designed to minimize sedation by limiting brain exposure, and loperamide, an antidiarrheal opioid, has very limited BBB permeability under normal conditions (Hu et al., 2015; Zoghbi et al., 2008). Unfortunately, many potential drug candidates fail late in development due to inadequate BBB penetration (Pardridge, 2005; Abbott et al., 2010), resulting in substantial financial costs and prolonged development timelines. Therefore, predicting BBB permeability early in the drug discovery process can drive candidate selection effectively, enabling more efficient allocation of experimental resources and reducing late-stage failures by identifying compounds with a higher likelihood of achieving effective CNS exposure (Kola & Landis, 2004; DiMasi et al., 2016). Here, it is important to consider that in some therapeutic contexts, limited BBB permeability is actually a desirable pharmacological property. Exposure of the CNS to certain drugs, such as second-generation antihistamines like loratadine and fexofenadine, can lead to unwanted side effects, and they are engineered to remain peripherally restricted (Kay, 2000; Simons & Simons, 2011; Rankovic, 2015). Traditionally, this prediction process has been attempted through experimental assays and physicochemical heuristics such as lipophilicity, molecular weight, and polar surface area (Bart et al., 2003; Carpenter et al., 2014; Di et al., 2003; Kaisar et al., 2016). However, these approaches are often resource-intensive and limited in predictive power (Carpenter et al., 2014).

Recent computational approaches have supported BBB permeability prediction through physicochemical descriptors, quantitative structure-activity relationship models, molecular fingerprints, and learned molecular representations (Yuan et al., 2018; Plisson & Piggott, 2019; Jafarpour et al., 2024; Pawar *et al.,* 2024). Early approaches based on physicochemical heuristics, such as lipophilicity, polar surface area, and hydrogen bonding, have gradually been complemented by quantitative structure-activity relationship (QSAR) models and, more recently, by machine and deep learning frameworks capable of learning directly from molecular graphs (Keep et al., 2020; Awe *et al.,* 2025; Enejoh *et al.,* 2025; Lamptey *et al.,* 2026; Babajide *et al.,* 2026). Public BBB data used in this work were obtained from the curated B3DB resource (Meng et al., 2021) and the MoleculeNet BBBP collection derived from the study by Martins et al. (2012). To assess the contribution of explicit three-dimensional and quantum-chemistry information under controlled conditions, we compared PaDEL plus Morgan features, the same features augmented with GFN2-xTB descriptors, and frozen Uni-Mol v1 representations using the same molecules and scaffold-based validation splits (Bannwarth et al., 2019; Zhou et al., 2023; Morgan, 1965).

Here, we propose BrainRoute, an open-source, community-driven platform for BBB permeability prediction and data curation. BrainRoute integrates heterogeneous BBB permeability datasets into a unified resource and provides a structured framework for accessing, verifying, and contributing experimental data. Rather than functioning as a static dataset or prediction tool, the platform is the primary contribution of this work and is designed to provide an evolving data infrastructure, where users can retrieve the relevant data and contribute to the growing dataset through the established verification and review pipeline. We aim to provide support for informed data selection and integration into the current dataset to enable future development of increasingly accurate and generalizable models.

## 2. Methodology

### 2.1 Data Collection and Integration

The data used for model development were obtained from two publicly available resources: B3DB and the MoleculeNet BBBP dataset (https://moleculenet.org/datasets-1). Both provide molecular structures and binary BBB annotations that can be redistributed and linked at the structure level. B3DB provides its published BBB+/BBB-classification, source reference codes, evidence-quality groups, numerical logBB values where available, and source-specific thresholds for a subset of records (Meng et al., 2021). For the categorical dataset, Group A records have numerical logBB values and B3DB applies a logBB threshold of −1; Group B records come from sources using the same −1 threshold and all sources agree on the categorical label; Group C records have agreement across sources but no reported threshold; and Group D records contain conflicting categorical reports for which B3DB retains the most prevalent label. B3DB discarded equal-frequency Group D ties before distributing the 7,807-record dataset. MoleculeNet BBBP provides the binary p_np endpoint derived from Martins et al. (2012); p_np = 1 was mapped to BBB+ and p_np = 0 to BBB-. Its distributed file does not contain row-level assay type, species, or numerical permeability values. These endpoints combine heterogeneous in vivo, in vitro, and literature-derived evidence and are not treated as one uniform assay. Figure 1 shows the overall workflow for platform development. Figures 2 and 3 show insights from the data collected through exploratory data analysis and representation.

**Figure 1.**
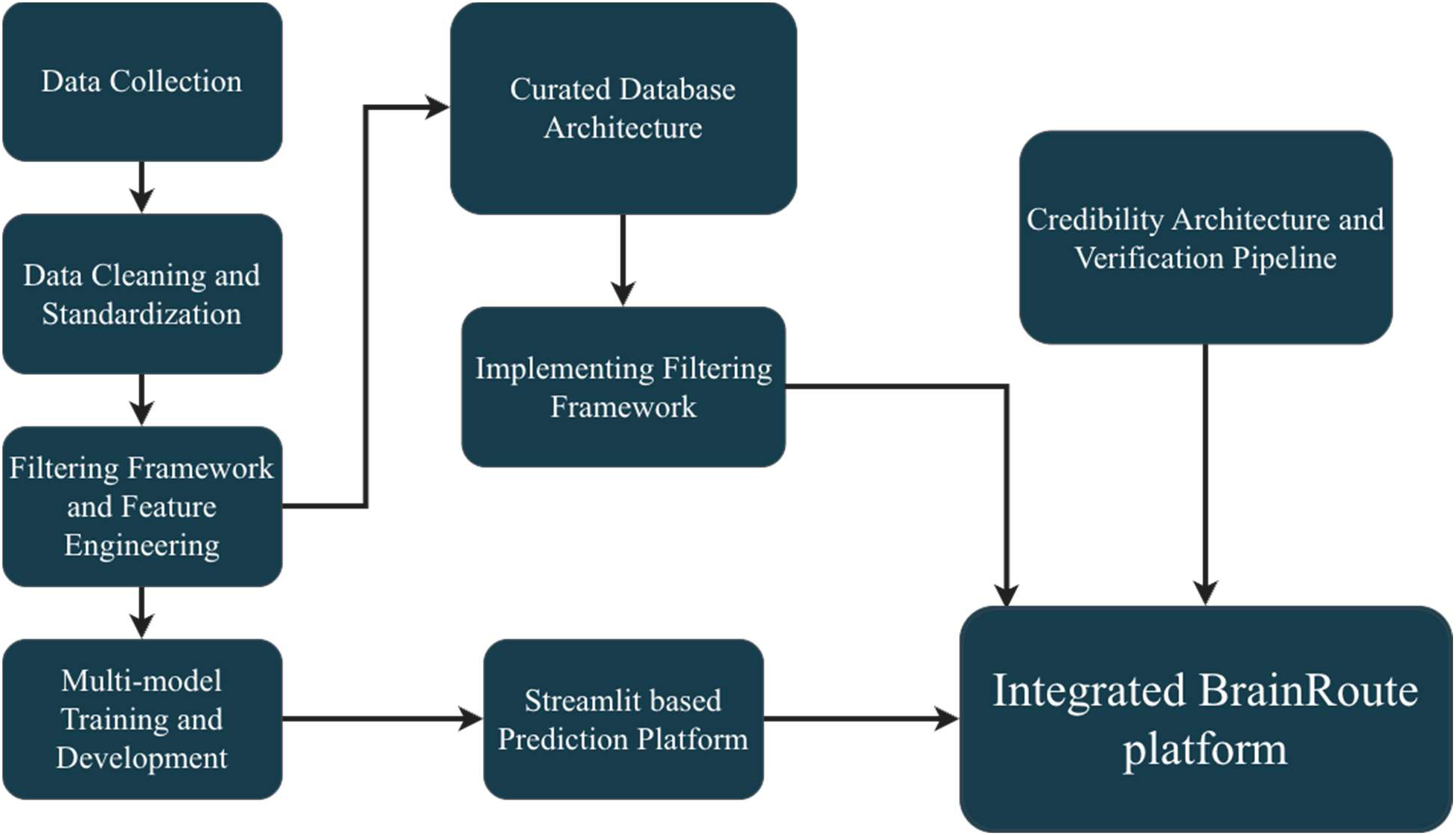
Overall computational workflow of the BrainRoute platform

**Figure 2.**
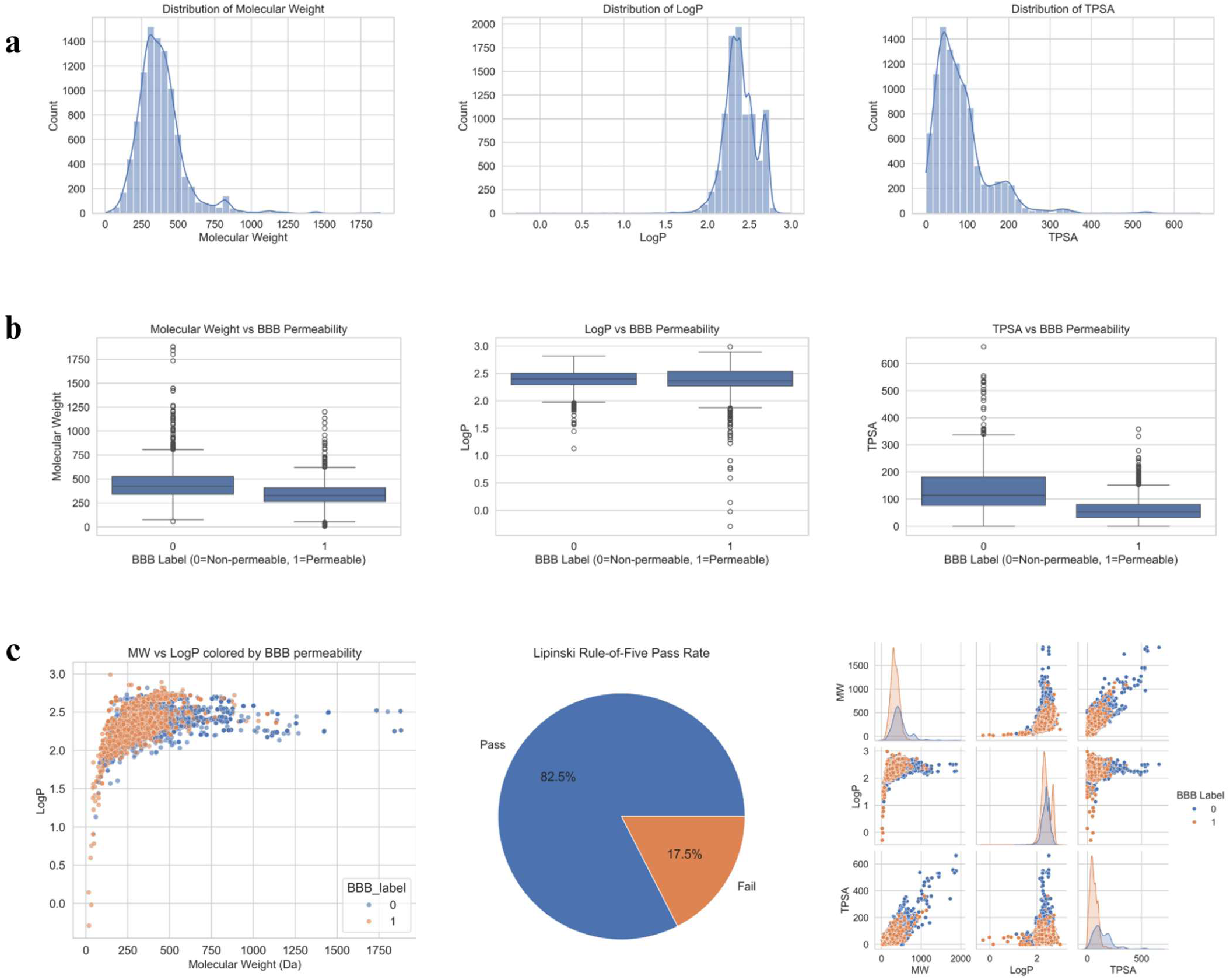
**a.** Distribution of Molecular Weight, LogP (measure of lipophilicity), and TPSA (Topological polar surface area). **b.** BBB permeable/non-permeable distribution among molecule descriptors (Molecular Weight, LogP, TPSA). **c.** 1-Permeability labeled distribution of molecules among descriptors (Molecular Weight, LogP), 2-Lipinski rule-of-five distribution of data, 3-Pairwise relationships of molecular descriptors

**Figure 3.**
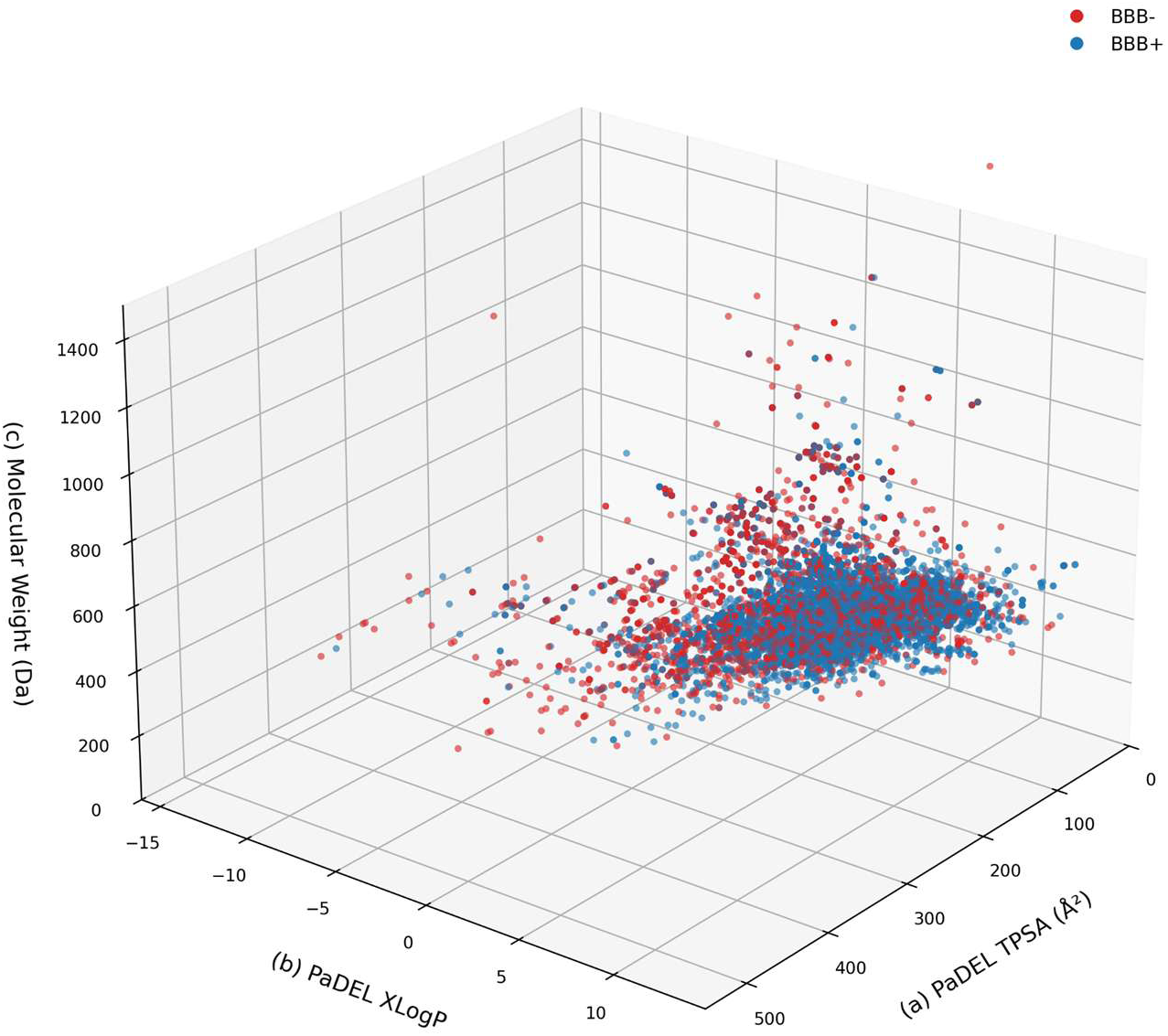
Representation of BBB molecule data in selected PaDEL Property Space with the following axes: **(a)** TPSA (Topological Polar Surface Area), **(b)** LogP (Lipophilicity), **(c)** Molecular Weight (Da). Each point represents one molecule, colored by BBB permeability class, with BBB−compounds shown in red and BBB+ compounds shown in blue.

The available B3DB file contained 7,807 records, including 4,956 BBB+ and 2,851 BBB-records. The available MoleculeNet BBBP file contained 2,050 records, including 1,567 BBB+ and 483 BBB-records. Together, they contained 9,857 source records. The repository records the source citation, file record count, SHA-256 file checksum, zero-based source row, source label, standardized structure, and InChIKey. References to the original datasets identify their published sources, while file checksums document the exact files used and row-level mappings link each modeled molecule to its original source record. These three forms of documentation serve different purposes and none is presented as a replacement for the others. B3DB and MoleculeNet BBBP were selected because they provide public structure-level binary BBB labels and support redistribution and row-level linkage. Other resources discussed in the Introduction were not added when their endpoints, access conditions, or overlap could not be harmonized into the same auditable binary task. This restriction limits coverage but avoids presenting unlike endpoints as directly interchangeable.

### 2.2 Data Cleaning and Standardization

For the original model cohort, RDKit was used to parse and sanitize the supplied structures, generate canonical isomeric SMILES and InChIKeys, identify exact structure duplicates, and calculate Bemis-Murcko scaffolds (Bemis and Murcko, 1996; Heller et al., 2013; RDKit, 2026). The original workflow did not apply an independent salt-stripping step, charge neutralization, tautomer canonicalization, or stereoisomer collapse. Disconnected components and formal charges were therefore retained in the original model representation, stereochemistry was retained in canonical isomeric SMILES and chirality-aware Morgan fingerprints, and Murcko scaffold grouping ignored chirality. Legacy PaDEL descriptors were calculated from the assembled input SMILES and then aligned to the standardized identities; Morgan and ChemBERTa representations were generated from the standardized cohort.

Molecular descriptors were computed using PaDELPy (Ecrl, 2019), a Python wrapper for PaDEL-Descriptor (Yap, 2011). PaDEL produced 1,875 numerical descriptors. The nAcid field was excluded to maintain compatibility with the original model artifacts, leaving 1,874 PaDEL inputs for modeling. Morgan fingerprints and frozen ChemBERTa-77M-MLM embeddings were also evaluated as molecular representations (Chithrananda et al., 2020; Rogers and Hahn, 2010).

Initial pipeline development used approximately 200 RDKit descriptors as a baseline, after which PaDEL descriptors were used to provide a larger set of physicochemical, topological, and geometric variables. The retained model matrix contains 1,874 of the 1,875 numerical PaDEL descriptors after the nAcid compatibility exclusion. The options provided by PaDELPy do not establish a clear conformer workflow. For this, the controlled GFN2-xTB and Uni-Mol benchmark therefore used an explicit and independently recorded conformer-generation protocol, as described below.

We also used Morgan fingerprints (nBits = 2048, useChirality = True) to add additional information and fixed-length molecular representation, suitable for ML model building. The 2048-bit ECFP4-like representations captured environments up to two bonds away from each atom (radius = 2) in a given molecule, giving important information about molecular similarity and preserving stereochemical information. (Rogers & Hahn, 2010)

The ChemBERTa-77M-MLM is a transformer-based masked language model trained on SMILES strings. These SMILES strings were tokenized and passed through the model with a maximum length of 256, aiming to keep the runtime manageable, as the dataset consisted of small molecules. Batch size of 32 and CLS pooling were used, creating a 384-dimensional (determined by the model’s hidden size) continuous embedding vector for each molecule. These embeddings were used to complement the rest of the feature space and efficiently capture SMILES-derived molecular representation information.

Using these molecular representations, we constructed multiple model-ready feature views to evaluate the contribution of each descriptor type individually and in combination. These included PaDEL descriptors alone, Morgan fingerprints alone, ChemBERTa embeddings alone, PaDEL + Morgan, and PaDEL + Morgan + ChemBERTa embeddings. We removed metadata columns such as SMILES, InChIKey, scaffold, molecule name, source information, and labels from the input feature matrices before training models to address feature leakage concerns, reduce the risk of metadata leakage, and restrict model inputs to molecular feature representations.

Figure 4 explains the data flow through the preprocessing and standardization steps to train models for the BrainRoute prediction tool. The two available source files contained 9,857 records. Independent source reconstruction retained 9,844 RDKit-valid records after 13 invalid structures and represented 8,090 unique InChIKeys. Across the source records, 48 unique InChIKeys had more than one BBB label. The legacy merged modeling table had separately identified and excluded 32 conflicting-label structures, producing a final original cohort of 7,888 molecules, including 5,424 BBB+ and 2,464 BBB-. Row-level source reconstruction identified 24 retained molecules, 0.30% of the original cohort, for which the available source records disagree. The original deployed artifacts are preserved for reproducibility, but all 24 molecules were excluded from the controlled benchmark. The original cohort contains Morgan and ChemBERTa representations for 7,888 molecules and PaDEL-aligned representations for 7,882. All curated molecules are available through the BrainRoute platform to help users to access, filter, and download any molecular records that they might need for further model development or data exploration.

**Figure 4.**
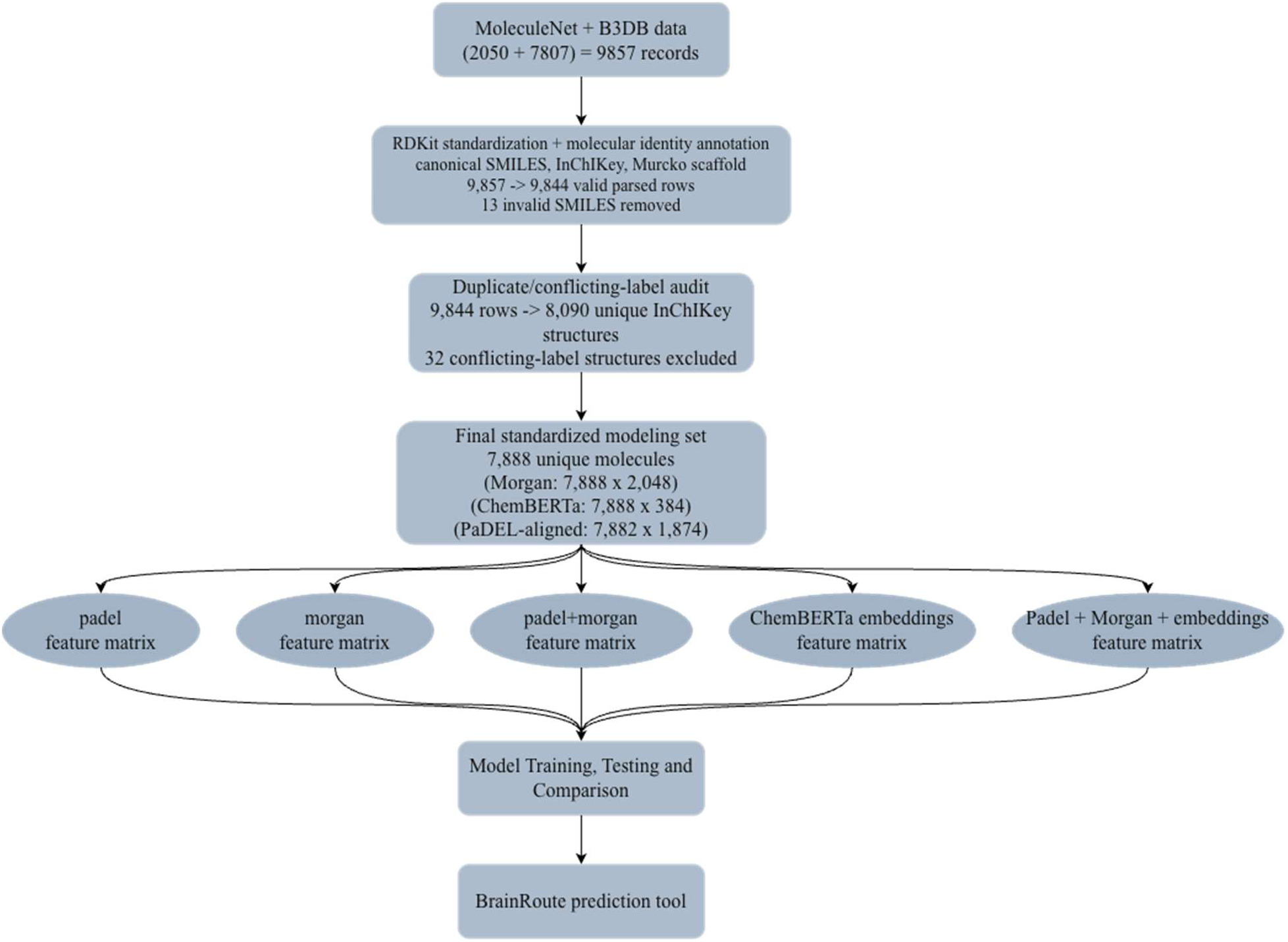
BrainRoute data curation and model development workflow. Data acquired from MoleculeNet and B3DB were standardized, audited for duplicates and conflicting labels, converted into molecular feature matrices, and used for BBB permeability model training, validation, and deployment.

### 2.3 Molecular Classification and Filtering Framework

To provide easy access to relevant data, we implemented a fast filtering and search system on the platform. This allows users to query molecules based on physicochemical properties, structural features, and drug-likeness rules. Physicochemical filters are based on descriptors that are known to influence molecular permeability and pharmacokinetics, such as Topological Polar Surface Area (TPSA) and LogP (a measure of lipophilicity). Structural filters were used to capture information about important aspects of molecular architecture, such as molecular size, aromaticity (aromatic vs non-aromatic), ring count, and presence of heterocycles. Furthermore, we also calculated higher-level classifications, such as whether molecules had peptide-like and lipid-like characteristics, to support domain-specific data inquiry. It is important to note here that these classifications are heuristic approximations based on the molecular descriptors we calculated and do not serve as strict biochemical definitions. Molecular weight was added to the platform as a filter due to its established effects on absorption, distribution, and permeability (Lipinski et al., 2001).

Furthermore, we calculated the following drug-likeness rules to aid fast filtering and data analysis within the platform - Lipinski’s rule of five (Molecular weight ≤ 500 Da, LogP ≤ 5, hydrogen bond donors (HBD) ≤ 5, and hydrogen bond acceptors (HBA) ≤ 10 (Lipinski et al., 2001; Gakpey et al., 2025; Ogbodo et al., 2023), Veber’s rule (TPSA ≤ 140 Å² and ≤ 10 rotatable bonds (Veber et al., 2002)), Egan’s rule (LogP ≤ 5.88 and TPSA ≤ 131 Å² (Egan et al., 2000)), and Ghose filter (160-180 Da molecular weight, −0.4 to 5.6 LogP, 40 to 130 molar refractivity (Ghose et al., 1999)). Here, a subset of the original Ghose filter was implemented due to descriptor constraints. The platform provides options to filter the data through binned ranges for these variables or to apply a strict pass/fail indicator to filter drug-likeness properties directly.

To support structure-based filtering, the platform also provides a limited structural-alert screen informed by the PAINS concept. The alerts are implemented in RDKit using SMARTS (SMiles ARbitrary Target Specification), a notation used to define molecular substructure patterns (Daylight Chemical Information Systems, Inc., n.d.; RDKit, 2026). Each molecule is compared with six custom SMARTS queries representing broad thiazole-like, catechol, quinone, Michael acceptor, cyanamide, and acrylamide motifs. A molecule is flagged when RDKit detects at least one matching arrangement of atoms and bonds. These queries provide a reduced structural screen and are not equivalent to the complete PAINS filters described by Baell and Holloway (2010). It is important to note here that an alert only indicates the presence of one of the selected substructures and is not evidence that the molecule is inactive, produces assay interference, or is experimentally problematic.

The CNS MultiParameter Optimization (MPO) score was calculated for the training molecules and is also calculated for new molecules submitted through the prediction tool. This score is used to estimate how favorable a compound’s physicochemical profile may be for CNS drug development (Wager et al., 2010). BrainRoute uses an adapted formulation calculated as the sum of six desirability terms: approximate CNS MPO = DlogP + DlogD + DMW + DTPSA + DHBD + DpKa. Each term ranges from 0 to 1, giving a total score between 0 and 6. The logP term has a value of 1 at or below 3 and decreases linearly to 0 at 5. The logD term has a value of 1 at or below 2 and decreases linearly to 0 at 4. The molecular-weight term has a value of 1 at or below 360 Da and decreases linearly to 0 at 500 Da. The hydrogen-bond-donor term has a value of 1 at or below 0.5 and decreases linearly to 0 at 3.5. The basic pKa term has a value of 1 at or below 8 and decreases linearly to 0 at 10. The TPSA term is 0 at 20 Å² or below, increases linearly to 1 between 20 and 40 Å², remains at 1 between 40 and 90 Å², and decreases linearly to 0 between 90 and 120 Å².

When ChemAxon calculations are available, the platform uses logD at pH 7.4 and the calculated basic pKa. When these values are unavailable, logP is used as a substitute for logD and a default pKa value of 8.0 is used. These substitutions, particularly the default pKa value, can shift the calculated score toward a more favorable value. It is therefore important to note that BrainRoute’s implementation should be interpreted as an approximate molecular property-based screening aid and not as the original validated CNS MPO score or a replacement for a full pharmacokinetic evaluation.

### 2.4 Platform Architecture

The platform uses a simple frontend-backend system where the molecular data is saved in a database using Supabase. The database consists of all the important physicochemical and structural descriptors needed to filter and showcase important information about the molecules. The platform is built using Next.js 14, React, TypeScript, Tailwind CSS, Supabase, and Recharts.

The current design of the platform involves a navigation box on the home screen that can be used to access all the available options and services offered. Users can submit and track verifications through the home page and launch the prediction tool to predict molecule BBB permeability. The platform allows users to sign in and track their verification submissions, predictions, and downloads through Supabase Auth. BrainRoute supports Google OAuth-based sign-in and passwordless magic link sign-in, which allows users to authenticate using established identity provider technologies (Google, 2026; Hardt, 2012; OpenID Foundation, 2014; Supabase, n.d.; Supabase, n.d.). This allows the platform to support user-specific record management and does not restrict access to the public BBB dataset.

Verification submissions require the users to be signed in so that submissions can be tracked and linked to the submitting account and user profile. Users can choose whether their verification submissions should be publicly visible or private. Public submissions are visible in the past submissions list to all the platform users. Private submissions are only visible to the submitting user through their authenticated profile. This design allows BrainRoute to support community-driven data submission and allows users to easily track their activity on the platform. The account-linked records are protected using Supabase Row-Level Security (RLS) policies. User records such as verification submissions, download history, and prediction tool interaction are scoped to the authenticated user, whereas public research records remain available for all users to interact with and download. This platform-level authorization model follows the principle of least privilege by separating public and private records based on the user identity (National Institute of Standards and Technology, n.d.; OWASP Foundation, n.d.-a; Supabase, n.d.; Supabase, n.d.). HTTPS/TLS handles the communication with authentication and database services, and Supabase reports encryption of hosted customer data at rest and in transit, providing an additional infrastructure-level security layer (OWASP Foundation, n.d.-b; Supabase, n.d.).

The backend for the database is also based on Supabase, which is used to host a PostgreSQL database, access policies, and object storage. The front end of the platform initializes Supabase using the public URL and then uses supabase-js to directly access table queries from the browser, making the implementation and access centralized and reusable. Because the Supabase anon key is exposed in the browser, database and storage access are controlled through Supabase Row Level Security policies rather than hidden server-side credentials.

The front end is designed like a scientific data exploration dashboard to better enable data analysis. The output region contains a live result count, a paginated data table, analytics cards and charts, and CSV export controls. Detailed molecular views provide comprehensive access to chemical information, including SMILES notation, physicochemical descriptors, structural properties, BBB/CNS permeability annotations, classification binnings, and adherence to drug-likeness rules. The verification page works like a form where users can submit information about the data they want to verify or add to the platform. The submissions then go through the credibility architecture described in section 2.6.

The prediction tool interface was developed using the Python Streamlit library, enabling rapid deployment of an interactive web application with custom CSS styling for improved usability and layout organization. Performance optimizations were implemented through caching and session state management, which reduced redundant computations and improved response times. The platform also supports export of prediction results and chat histories to facilitate offline documentation and analysis.

BrainRoute supports both single-molecule prediction and batch processing. For each molecule, the three deployed models produce BBB+ probabilities p1, p2, and p3. The ensemble probability is their arithmetic mean, p_bar = (p1 + p2 + p3)/3, and BBB+ is assigned when p_bar is at least 0.5. The interface field called confidence is 100 times max(p_bar, 1 - p_bar). The displayed uncertainty is 100 times the population standard deviation of p1, p2, and p3, and agreement is the percentage of individual thresholded labels equal to the ensemble label. These are descriptive ensemble scores, not formal posterior probabilities or guarantees of correctness. Internal scaffold out-of-fold and external ensemble Brier scores were 0.143 and 0.209, and the corresponding 10-bin expected calibration errors were 0.080 and 0.111. The weaker external calibration requires cautious interpretation of the interface confidence label. In batch mode, users can upload CSV files or paste molecular data directly, with predictions generated from individual models as well as an ensemble output based on averaged probabilities. The interface additionally provides visual and quantitative summaries, including class distribution charts and confidence–uncertainty plots, to aid interpretation. A file size limit of 200 MB is imposed due to inherent constraints of the Streamlit framework, which relies on in-memory data handling and server-side execution (Streamlit 2024). Larger uploads can lead to memory bottlenecks and degraded performance, particularly in web-deployed environments, making this limit necessary to maintain stability and responsiveness.

Permeability predictions made through user interaction with the prediction tool are directly added to the platform, with the predicted tag in the molecule card. Verified molecules have the verified tag, and molecules added to the database using the inbuilt prediction tool are given the predicted tag. Prediction confidence is also added to the molecule card of each molecule added to the database through the prediction tool. This system acts as an important layer to make the platform easy to interpret and user-friendly.

The Meta Llama 3 8B Instruct (Meta AI, 2024) model is currently incorporated in the BrainRoute prediction tool to help provide contextual interpretation of model outputs. The model has not been formally assessed for BBB-relevant mechanistic interpretations and is therefore included only as a supplementary aid for general understanding of prediction outputs, rather than a definitive scientific explanation. Users are advised to independently verify information generated by the chatbot. We have included a comparative evaluation of Llama 3 8B on various biomedical benchmarks in the GitHub repository.

### 2.5 Predictive Models Pipeline

For the BrainRoute prediction tool, we implemented a preprocessing, training, and model-selection pipeline. An ensemble approach was used so that the final prediction did not depend on a single classifier or molecular representation. LazyPredict (Pandala, 2023) was first used as an exploratory screen of approximately 30 supervised classification models. This screening considered accuracy, balanced accuracy, and F1 score and was used only to reduce the number of candidate model families. It is not treated as an independent estimate of final model performance. Based on this screening, six classifiers with different model structures were selected for further evaluation: K-Nearest Neighbors (KNN), Logistic Regression (LR), Random Forest (RF), Extra Trees (ET), Light Gradient Boosting Machine (LightGBM), and eXtreme Gradient Boosting (XGBoost) (Cover and Hart, 1967; Cramer, 2002; Breiman, 2001; Geurts et al., 2006; Ke et al., 2017; Chen and Guestrin, 2016; Hanson *et al.,* 2024).

The classifiers were evaluated using the five feature views described in Section 2.2: PaDEL descriptors, Morgan fingerprints, ChemBERTa embeddings, PaDEL plus Morgan, and PaDEL plus Morgan plus ChemBERTa. Following the development comparisons, three configurations were selected for the prediction tool: PaDEL plus Morgan LightGBM, PaDEL plus Morgan

Extra Trees, and PaDEL plus Morgan plus ChemBERTa XGBoost. These configurations were refitted on all 7,882 molecules with complete PaDEL representations before deployment. The three models remain the production ensemble used by BrainRoute, and their BBB+ probabilities are combined using the procedure described in Section 2.4.

Exact molecular duplicate handling was completed during data curation before any training and test partitions were created. Molecules were consolidated using standardized InChIKeys, and duplicate removal is therefore treated as a data-curation requirement rather than a validation strategy. A stratified random 80/20 split using seed 42 was retained as a conventional baseline. Five additional 80/20 molecule-level random hold-outs were generated using seeds 1, 2, 3, 4, and 5. Although these five runs grouped rows by InChIKey, every InChIKey was already unique at that stage. They are therefore described as secondary repeated random molecule-level hold-outs rather than duplicate-aware validation.

The primary internal evaluation used five-fold Bemis-Murcko scaffold-grouped cross-validation. A separate scaffold-group hold-out using seed 42 was included as a secondary analysis. In each scaffold-grouped split, molecules sharing the same core scaffold were assigned entirely to one partition. This provides a more demanding assessment of whether the models can generalize to compounds with core structures that were not represented in the corresponding training set. Split audits confirmed that there was no InChIKey overlap between training and test data for any evaluated split and no scaffold overlap within the scaffold-grouped analyses.

The curated modeling cohort contained 7,888 molecules, including 5,424 BBB+ and 2,464 BBB-molecules. This corresponds to a positive-to-negative ratio of 2.20:1, with BBB+ molecules representing 68.8% of the cohort. The PaDEL-complete subset used by the production models contained 5,422 BBB+ and 2,460 BBB-molecules. BBB+ is therefore the majority class in the training data. This direction of imbalance is important because curated BBB datasets are commonly enriched for BBB-permeable compounds and may not represent the prevalence expected in broader chemical screening populations (Martins et al., 2012; Huang et al., 2024).

Raw training, undersampling, SMOTE, and class-weighted training were considered during model development (Chawla et al., 2002; Meng et al., 2025; Fernández et al., 2018). Because the dataset is imbalanced toward BBB+ molecules, undersampling would have discarded a substantial number of majority-class compounds, and SMOTE would have required generating synthetic points in descriptor, fingerprint, or embedding space with no guarantee that these interpolated points correspond to chemically valid or physically realizable molecules. Class weighting avoids both problems as it preserves the full training set and lets each model penalize misclassification of the minority (BBB-) class more heavily during optimization, rather than altering the underlying data distribution.Hence, the production pipeline used model-specific class weighting. Logistic Regression, Random Forest, Extra Trees, and LightGBM used class_weight = “balanced.” XGBoost used scale_pos_weight equal to the number of BBB-molecules divided by the number of BBB+ molecules within each training partition. KNN does not provide a class-weight parameter, and uniform and distance-based neighbor weighting were included in its hyperparameter search.

Balanced accuracy was used as the primary model-selection metric because it averages sensitivity and specificity and therefore gives equal importance to performance in the BBB+ and BBB-classes, regardless of their different frequencies. Sensitivity and specificity were also reported separately to show whether a model favored the BBB+ majority class at the expense of identifying BBB-molecules. Precision, recall, and F1 score were retained as secondary measures because they describe positive-class prediction but depend on class prevalence and should not be interpreted alone. MCC was included as a summary of all four confusion-matrix outcomes, while AUROC assessed ranking performance across thresholds. AUPRC was interpreted relative to the BBB+ prevalence of each evaluation cohort because its baseline changes with positive-class frequency. The reasons for using these metrics are particularly important here because the internal and external cohorts have different class distributions.

During training, preprocessing and feature-filtering operations were fitted using only the training partition of each split and were then applied without refitting to the held-out molecules. Non-finite values were converted to missing values, variables with more than 15% missingness were removed, zero-variance variables were excluded, remaining missing values were replaced using training-set medians, and variables with an absolute Pearson correlation greater than 0.95 were filtered. StandardScaler was applied to LR and KNN. The missingness and correlation thresholds were fixed preprocessing rules and were not optimized against held-out performance (Kato et al., 2023; Kauffman and Jurs, 2001; Rácz et al., 2019; Trinh et al., 2023).

Model hyperparameters were selected using GridSearchCV with balanced accuracy as the scoring metric (Pedregosa et al., 2011). Within each held-out production-model split, the search used three-fold StratifiedKFold cross-validation applied only to the training partition. Scaffold grouping was used for the outer scaffold folds but was not applied to these inner production-model searches. The broader classifier-family and production-configuration screening was also not repeated within every outer scaffold fold. The internal production-model estimates may therefore contain some selection optimism and are interpreted as development estimates. The independent external validation set was not used for classifier screening, feature selection, hyperparameter tuning, production-configuration selection, or classification-threshold selection. The controlled benchmark described in Section 2.5.1 provides a stricter comparison in which scaffold grouping is applied to both the inner and outer validation loops.

As a leakage-control analysis, the BBB labels were randomly permuted using three fixed seeds. For each seed, the model pipeline was retrained using shuffled training labels and evaluated using the unchanged held-out labels. Balanced accuracy and AUROC decreased to approximately chance level across the tested models. This did not identify evidence of obvious label or metadata leakage, although label permutation does not replace scaffold-grouped or independent external validation (Ojala and Garriga, 2010; Tropsha, 2010).

A nearest-neighbor similarity audit was also performed using Morgan-fingerprint Tanimoto similarity (Rogers and Hahn, 2010; Willett et al., 1998). Each held-out molecule was compared with the corresponding training set, and its nearest-training similarity was recorded. The repeated random molecule-level hold-outs had a mean nearest-neighbor similarity of approximately 0.70, compared with approximately 0.50 for the scaffold-grouped folds. This supports the use of scaffold-grouped evaluation as the primary internal analysis because the random hold-outs contained more close structural analogues. The similarity audit was used for interpretation and did not affect model training.

#### 2.5.1 Controlled 3D and quantum-chemistry benchmark

A controlled matched benchmark compared three representations: the 3,922-feature PaDEL plus Morgan control, the same control augmented with 19 GFN2-xTB variables, and a frozen 512-dimensional Uni-Mol v1 molecular representation (Bannwarth *et al.,* 2019; Zhou *et al.,* 2023). Unlike the GFN2-xTB descriptors, which produce a small, fixed set of scalar features that can be concatenated directly onto the existing PaDEL plus Morgan matrix, the Uni-Mol v1 representation is a dense, 512-dimensional embedding produced by a separately pretrained SE(3)-equivariant transformer; we therefore evaluated it as a standalone representation rather than as an additive feature block, since naively concatenating a dense learned embedding with hand-crafted tabular descriptors would not let the classical tree-based classifiers weight the two feature types appropriately without a dedicated fusion architecture. Combining Uni-Mol embeddings with the existing descriptor set through a proper fusion or gating layer is a natural extension for future, GPU-supported iterations of this benchmark.

GFN2-xTB variables included total and electronic energy, HOMO and LUMO energies, HOMO-LUMO gap, dipole magnitude, isotropic polarizability, atomic-charge summaries, atomic-dipole summaries, Wiberg bond-order summaries, electron count, and unpaired-electron count. For each xTB calculation, the largest organic fragment was selected only for the 3D calculation, formal charge and stereochemistry were retained, one ETKDGv3 conformer was generated with a deterministic seed, and MMFF94s minimization with UFF fallback preceded a gas-phase GFN2-xTB single point.

Uni-Mol v1 used the official pretrained molecular weights through unimol_tools 0.1.6. RDKit hydrogen-added coordinates were generated with seed 42 and MMFF optimization where available. Representations produced from a 2D fallback were recorded and excluded. The benchmark also excluded xTB failures and the 24 retained source-label conflicts from all three feature views. The resulting matched cohort contained 7,740 internal molecules (5,310 BBB+ and 2,430 BBB-) and 410 external molecules (158 BBB+ and 252 BBB-). No representation received additional molecules. XGBoost was used for all three views with the same fixed hyperparameter grid. The outer five scaffold folds estimated performance; each outer training partition contained a three-fold StratifiedGroupKFold inner search grouped by scaffold. The independent external probabilities were averaged across the five fitted outer models. All thresholded metrics used 0.5.

#### 2.5.2 Alternative approaches tested

To further test out different approaches, we also trained models with just BBB-relevant descriptors. We chose this subset of PaDEL descriptors to capture important physicochemical and structural properties that affect passive diffusion across the BBB, including lipophilicity (e.g., ALogP, XLogP, MLogP), molecular size and weight (MW, AMW), polarity and surface area (TopoPSA, TPSA, PPSA/DPSA variants), hydrogen bonding capacity (nHBDon, nHBAcc), and molecular flexibility and topology (nRotB, ring counts, Zagreb index). These descriptors were selected by considering well-established principles in CNS drug discovery and design, where optimal BBB permeability is usually associated with moderate lipophilicity, low polar surface area, limited hydrogen bonding capacity, and controlled molecular size and flexibility (Pajouhesh & Lenz, 2005; van de Waterbeemd et al., 1998; Kelder et al., 1999; Lipinski et al., 2001). These models were not included in the prediction tool as their performance was worse compared to the models we trained using all other descriptors.

We also developed a deep learning approach, using the Structure-Property Multi-Modal (SPMM) foundational model developed by Chang, J. et al. The model architecture consists of a pre-trained SMILES encoder, which is based on the BERT (Bidirectional Encoder Representation from Transformers) architecture (Salah *et al.,* 2026). It treats the SMILES input strings as textual sequences and runs them through a self-attention mechanism that allows the model to learn contextual relationships between each token (atom, bond, or substructure fragment). This enables the encoder to capture how the local environment of a substructure (e.g., a nitrogen atom adjacent to an aromatic ring) influences its overall molecular behavior. The training and evaluation workflow was implemented in PyTorch, incorporating the Hugging Face Transformers library for tokenization and BERT weight handling. We did not connect the trained model to the main platform as the package versions required for the SPMM scripts conflict with the current BrainRoute software stack. This represents a compatibility limitation within the current BrainRoute platform architecture rather than a limitation of the SPMM model itself. We do not intend to integrate this model in future versions of the platform as it would require significant changes to the backend workflow or a separate isolated deployment workflow, which is out of the scope of the present release. However, deep learning approaches remain within the broader future scope of BrainRoute if they can be implemented using compatible dependencies or a separate model-serving framework.

#### 2.5.3 Testing and Deployment

To validate our models, we used several standard classification metrics, such as:

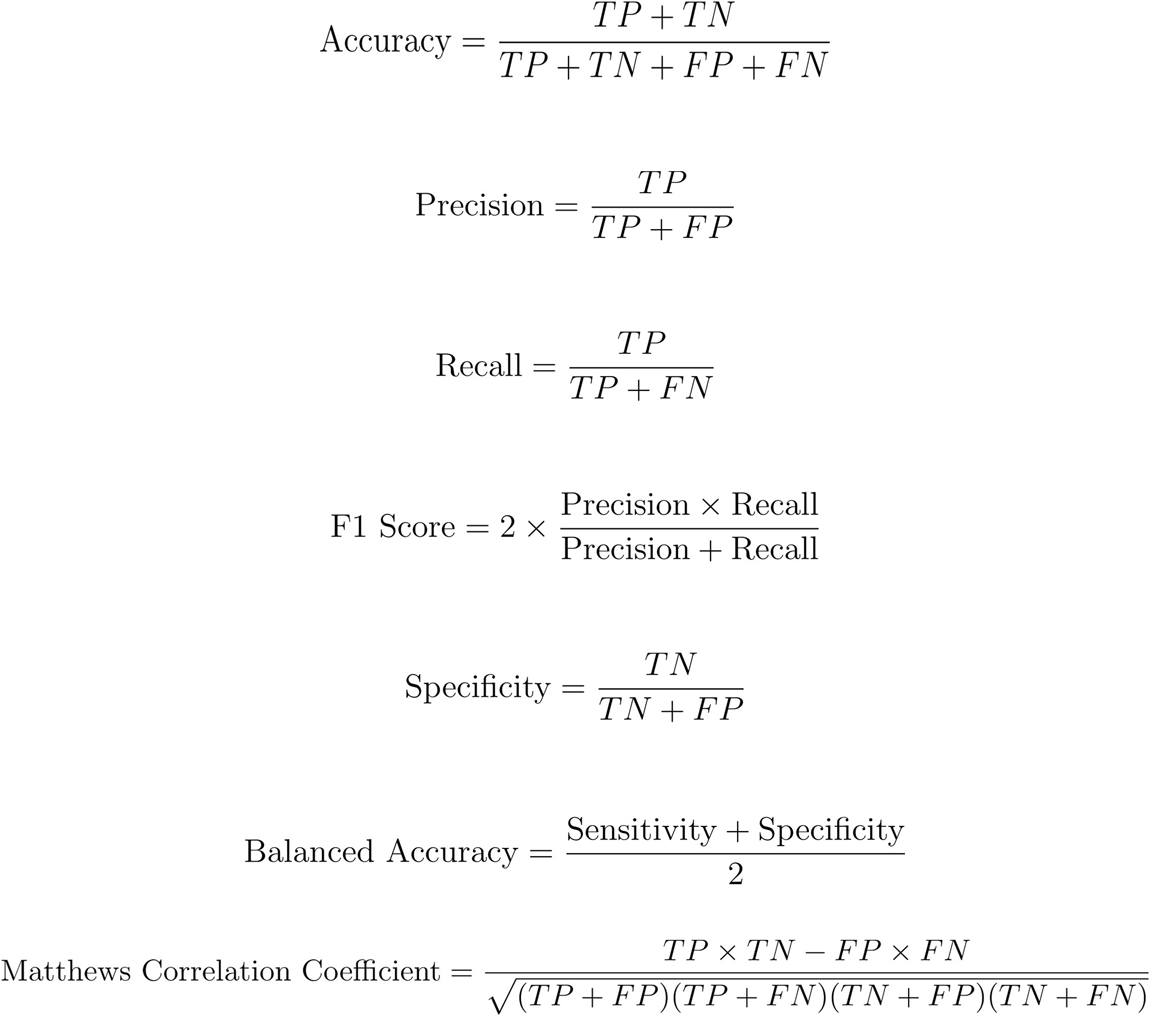

where TP = True Positives, TN = True Negatives, FP = False Positives, FN = False Negatives

Model performance was evaluated using AUROC, AUPRC, balanced accuracy, sensitivity, specificity, precision, negative predictive value, F1, and MCC. Because BBB+ is the positive class and is the majority class, AUPRC is interpreted relative to positive prevalence and is not used alone. Probability quality was assessed using Brier score and 10-bin expected calibration error. A fixed threshold of 0.5, selected before evaluation, converted predicted probabilities to BBB+ or BBB-labels; no held-out test set was used to tune the threshold.

Outer scaffold-fold performance is reported as mean plus or minus standard deviation. For pooled out-of-fold and external predictions, 95% confidence intervals were calculated using 2,000 bootstrap resamples clustered by Bemis-Murcko scaffold. Paired bootstrap differences and exact McNemar tests were used for matched predictions, with Holm adjustment across the planned comparisons. Historical five-fold Wilcoxon tests are reported with exact two-sided p-values, but five-fold pairs have limited power and a non-significant result is not interpreted as evidence of model equivalence.

Balanced accuracy was the primary selection metric because the original cohort contains 5,424 BBB+ molecules and 2,464 BBB-molecules, a positive-to-negative ratio of 2.20:1. The positive class therefore represents 68.8% of the cohort. Sensitivity and specificity are reported separately so performance in both classes remains visible. All additional calculations and training were performed on CPU-based Apple silicon with 16 GB unified memory; no GPU or paid quantum-chemistry service was used.

#### 2.5.4 Independent external validation

The deployed models were evaluated on an independent external validation set (IVS) obtained from the BBB dataset reported by Faramarzi *et al*. (2022). This dataset was not used for classifier-family screening, preprocessing-rule selection, feature-configuration selection, hyperparameter tuning, production-model selection, or classification-threshold selection.

The retrieved external dataset contained 921 records. Before evaluation, 358 molecules with an exact InChIKey overlap with the internal modeling cohort were removed. Complete PaDEL descriptors could not be generated for a further 153 molecules. The final IVS therefore contained 410 non-overlapping, PaDEL-complete molecules, including 158 BBB+ and 252 BBB-molecules. The BBB+ prevalence in the IVS was 38.5%, compared with 68.8% in the internal cohort.

Each of the three full-data production models was evaluated separately on the IVS, followed by evaluation of their probability-averaged ensemble. The included molecules, exact-overlap exclusions, and PaDEL-failure exclusions were compared using class distribution, molecular weight, logP, TPSA, nearest-training Tanimoto similarity, and scaffold overlap. This analysis showed that external-set attrition was not random, particularly for molecules that did not produce complete PaDEL output. The external results therefore describe performance on the 410 molecules that passed identity screening and feature generation and should not be interpreted as an unbiased estimate for every possible chemical structure. Individual model results, ensemble results, class-specific metrics, and confidence intervals are reported in the Results and supplementary material.

### 2.6 Credibility Architecture and Verification Pipeline

Through BrainRoute, we aim to create a community-driven, evidence-backed molecular database where the data can be traced back to verifiable literature sources. For this, we designed and implemented a multi-stage validation process before influencing the canonical dataset. These submissions are not restricted to the original authors of the papers, provided the user supplies complete, verifiable experimental data and a valid DOI, along with other requirements outlined below. At the current version of the platform, the validation process is conducted manually to ensure careful, context-aware interpretation of experimental results submitted. This approach allows us to establish a structured foundation for future scalability.

#### User submission workflow

Users submit information through the “Verify Data” page of the platform. On this page, they can submit user information, a valid DOI, the technique used, experiment description, experimental results/data, permeability indication, and supporting files for the submission. Users must be signed in to their BrainRoute account to submit a verification request. This page allows users to submit information about whether the study was in vitro, in vivo, or in silico, the specific assay used, the cell line or organism involved, and all other relevant experimental factors such as concentration ranges or detection methods used. This allows users to provide experimental context for each submission rather than an isolated label. User verification submissions are not automatically added to the database and are reviewed through our verification process first. Verification submissions with insufficient information provided would not be considered for inclusion. Predictions generated through the BrainRoute prediction tool are added to the overall dataset with a “br_predicted” tag. These model-derived entries remain as distinguishable records within the browsable dataset and are not treated as experimentally verified labels unless they are subsequently supported through the verification workflow. This allows users to include or exclude predicted entries during downstream filtering, export, or model development.

#### Verification workflow

All submissions are routed through a manual verification process conducted by the BrainRoute development team before the data can be integrated into the database. In this stage, the given metadata with the submission will be used to confirm the reported values and interpretations of experimental outcomes, along with the accuracy of reported methods and the background of the study. This process allows for nuanced judgment in certain cases where experimental design or reporting can introduce ambiguity.

#### Tiering System and Tracking

We have implemented a tier system to account for differences in the experimental reliability of each user submission, which reflects the strength of the underlying data. High-confidence entries correspond to well-characterized experimental studies with complete methodological reporting, particularly those involving in vivo validation or widely accepted in vitro systems. Moderate-confidence entries include studies with partial metadata or limited validation, while computational predictions are categorized separately to distinguish model-derived outputs from experimental evidence. Preliminary or incomplete submissions are retained with appropriate labeling to indicate the need for further verification. This tiering framework allows the platform to integrate diverse forms of evidence while preserving transparency about their relative reliability and intended use.

The submissions are stored within a version-controlled framework that stores the history of submissions and any subsequent modifications. Users are able to edit their verification submissions through their profile page and can track the status of their submissions there as well. Once submitted, the submission status can be changed to any of the following options based on submission content: “Submitted”, “In Review”, “Accepted”, “Denied”, or “More information requested”. Verification submissions that are denied for inclusion in the dataset are removed from the active user-facing submission list after one week but retained in an internal audit log for provenance and administrative tracking. This structure allows the community to track individual data points over time, provides accountability for changes made within the system, and supports the clear communication needed for a transparent verification process.

Conflicting reports of expected BBB permeability features due to variability in experimental systems and measurement approaches are identified and preserved within the platform. Submissions are flagged with multiple studies suggesting different quantitative values for the same molecules. Each case is then evaluated in the context of experimental design, with consideration given to assay type, biological model, and overall methodological rigor. When one source provides clearly stronger evidence, it is emphasized while still retaining all associated records. If the verification data changes provided by users clearly and accurately show a different outcome than a model prediction, the molecule BBB value and tag can be changed accordingly. In cases where multiple sources are of comparable quality for the same molecule, a “Conflicting Evidence” tag is added for the molecule, and all underlying studies remain accessible. Molecules are also tagged as “br_training”, “br_verified”, and “br_predicted”, based on whether they were used for the BrainRoute prediction tool model training, verified through the verification system, or predicted using the prediction tool, respectively. All information about the molecules is available in the individual molecule card and can be downloaded through the platform. This approach ensures that users are presented with the full landscape of available data and can interpret discrepancies in an informed manner.

## 3. Results

The highest-performing feature configuration for each model type is reported in Table 1. Two internal validation settings were used to provide different estimates of model performance: five repeated random 80/20 molecule-level hold-outs and five-fold Bemis-Murcko scaffold-grouped cross-validation. Exact molecular duplicates had already been consolidated by standardized InChIKey before either validation procedure was performed. The repeated random hold-outs therefore do not represent a separate duplicate-control strategy. Instead, they provide a conventional estimate of performance when the training and test sets may contain structurally related molecules. In contrast, the scaffold-grouped analysis assigned all molecules sharing the same core scaffold to a single fold, requiring the models to predict molecules with core structures that were not represented in the corresponding training data.

**Table 1.** Performance summary for best-performing models across all feature views and validation strategies.

| Model | Feature view | Balanced accuracy | AUPRC | MCC | F1 | AUROC |
|---|---|---|---|---|---|---|
| **LightGBM** | PaDEL + Morgan | 0.836 ± 0.008 | 0.959 ± 0.002 | 0.657 ± 0.016 | 0.886 ± 0.006 | 0.916 ± 0.004 |
| **Extra Trees** | PaDEL + Morgan | 0.830 ± 0.011 | 0.957 ± 0.003 | 0.636 ± 0.019 | 0.874 ± 0.007 | 0.905 ± 0.005 |
| **XGBoost** | PaDEL + Morgan + Embeddings | 0.827 ± 0.012 | 0.952 ± 0.004 | 0.631 ± 0.024 | 0.872 ± 0.009 | 0.902 ± 0.008 |
| **Random Forest** | Morgan | 0.821 ± 0.013 | 0.958 ± 0.003 | 0.629 ± 0.019 | 0.877 ± 0.007 | 0.911 ± 0.003 |
| **Logistic Regression** | PaDEL | 0.786 ± 0.002 | 0.923 ± 0.004 | 0.543 ± 0.004 | 0.831 ± 0.005 | 0.849 ± 0.004 |
| **KNN** | PaDEL + Morgan | 0.774 ± 0.013 | 0.908 ± 0.006 | 0.567 ± 0.024 | 0.870 ± 0.007 | 0.861 ± 0.010 |

Model performance was consistently higher under the repeated random molecule-level hold-outs than under scaffold-grouped cross-validation (Table 2). The highest mean balanced accuracy in the random hold-outs was obtained by the PaDEL plus Morgan LightGBM model, with a balanced accuracy of 0.836 ± 0.008. Under scaffold-grouped cross-validation, the highest mean balanced accuracy was obtained by the PaDEL plus Morgan plus ChemBERTa XGBoost model, with a balanced accuracy of 0.783 ± 0.026. The nearest-neighbor analysis showed a mean nearest-training Tanimoto similarity of approximately 0.70 for the random hold-outs and approximately 0.49 for the scaffold-grouped folds. This confirms that molecules in the random test sets were, on average, more similar to the corresponding training molecules. The lower scaffold-grouped performance is therefore consistent with the more difficult task of predicting molecules with less familiar structural frameworks, rather than with the presence or absence of exact duplicates. For this reason, the scaffold-grouped analysis is treated as the primary internal estimate of generalization, while the repeated random hold-outs are retained as a secondary and less stringent comparison. Neither internal analysis replaces the independent external validation reported below.

**Table 2.** Top three classification configurations within both validation strategies.

| **Validation strategy** | **Model** | **Feature view** | **Balanced accuracy** | **AUPRC** | **MCC** | **F1** |
|---|---|---|---|---|---|---|
| **Repeated random molecule-level hold-out** | LightGBM | PaDEL + Morgan | 0.836 ± 0.008 | 0.959 ± 0.002 | 0.657 ± 0.016 | 0.886 ± 0.006 |
| **Repeated random molecule-level hold-out** | Extra Trees | PaDEL + Morgan | 0.830 ± 0.011 | 0.957 ± 0.003 | 0.636 ± 0.019 | 0.874 ± 0.007 |
| **Repeated random molecule-level hold-out** | XGBoost | PaDEL + Morgan + Embeddings | 0.827 ± 0.012 | 0.952 ± 0.004 | 0.631 ± 0.024 | 0.872 ± 0.009 |
| **Scaffold-aware CV** | XGBoost | PaDEL + Morgan + Embeddings | 0.783 ± 0.026 | 0.931 ± 0.022 | 0.543 ± 0.047 | 0.839 ± 0.037 |
| **Scaffold-aware CV** | LightGBM | PaDEL + Morgan | 0.780 ± 0.034 | 0.934 ± 0.020 | 0.547 ± 0.062 | 0.851 ± 0.035 |
| **Scaffold-aware CV** | Extra Trees | Morgan | 0.778 ± 0.023 | 0.936 ± 0.023 | 0.549 ± 0.054 | 0.847 ± 0.045 |

Furthermore, comparing feature views across both validation strategies revealed that the PaDEL+Morgan representation of data gave the highest balanced accuracy, AUPRC, MCC, and tied for the highest F1 score (Table 3). This shows that adding frozen ChemBERTa embeddings to the feature space did not substantially improve the average performance when utilized as fixed tabular features for classical ML models, as the PaDEL + Morgan + ChemBERTa embeddings feature space performed nearly identically to the PaDEL + Morgan feature space, while being slightly lower across balanced accuracy, AUPRC, and MCC. This suggests that the traditional physicochemical descriptors contributed significantly to the models’ internal representation as tested in this study. A detailed table with all performance metrics for all training rounds is provided in the GitHub repository.

**Table 3.** Average performance by feature space across all validation strategies and models.

| **Feature space** | **Balanced accuracy** | **AUPRC** | **MCC** | **F1** |
|---|---|---|---|---|
| **PaDEL + Morgan** | 0.782 | 0.926 | 0.553 | 0.852 |
| **PaDEL + Morgan + Embeddings** | 0.781 | 0.925 | 0.552 | 0.852 |
| **PaDEL** | 0.773 | 0.922 | 0.529 | 0.837 |
| **Morgan** | 0.766 | 0.917 | 0.529 | 0.847 |
| **Embeddings** | 0.728 | 0.897 | 0.448 | 0.816 |

However, it is important to note that this finding should not be interpreted as a general limitation of the use of ChemBERTa for molecular modelling, as the frozen embeddings were passed into classical machine learning models and not trained directly for BBB permeability prediction.

Detailed metrics with all trials, including standard deviation information, fold-level metrics, pooled common-molecule predictions, confidence intervals, exact p-values, and split-overlap audits are provided in the GitHub repository. These numbers reflect a robust pipeline, highly reducing the risk of data leakage and sampling issues affecting the results. Figure 5 helps to further visualize performance metrics for these models. A baseline value of the AUPRC curve (Figure 6), representing the prevalence of the positive data class, was 68%.

**Figure 5.**
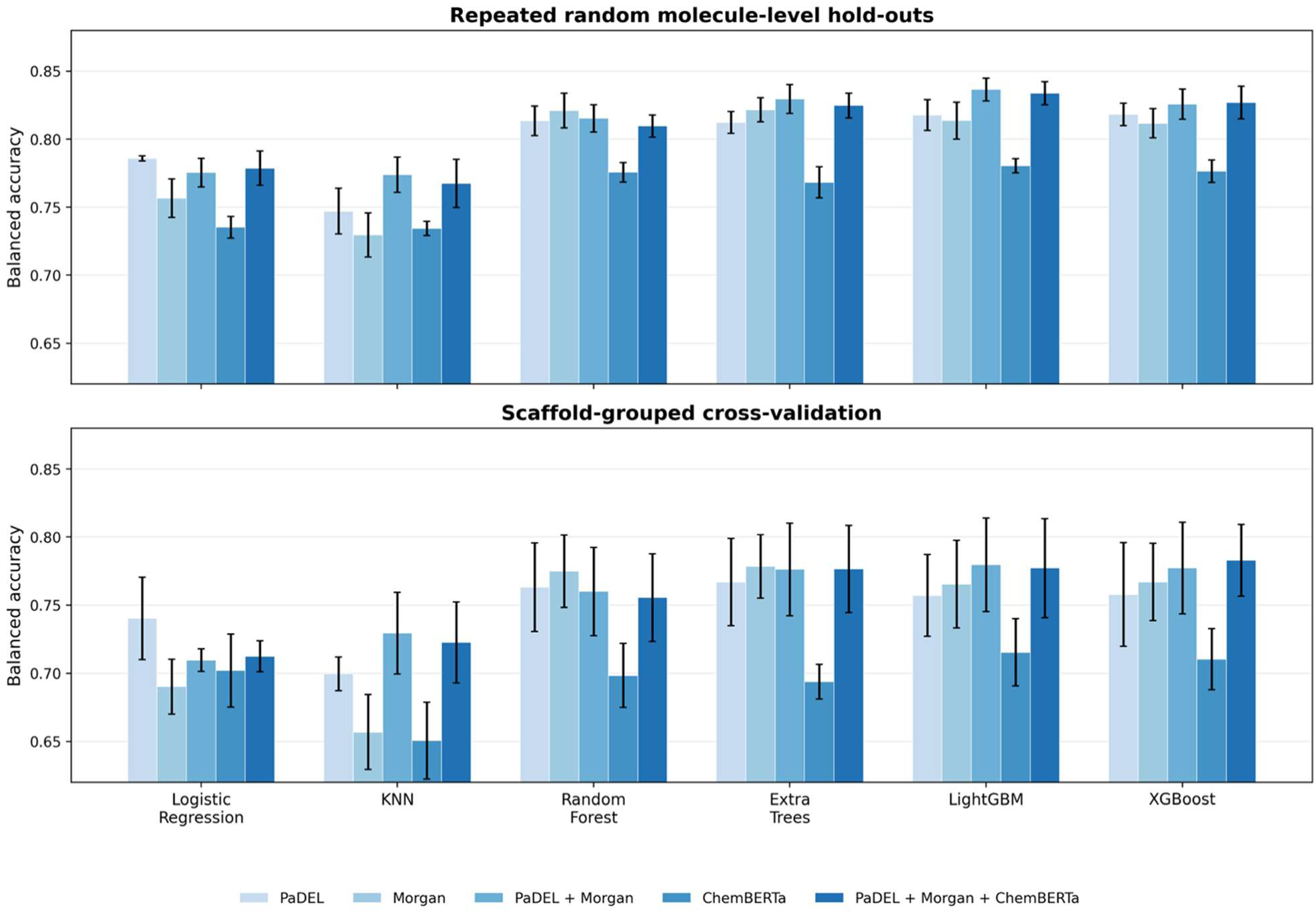
Balanced accuracy performance for all models and feature spaces for both - Repeated random molecule-level hold-out and Scaffold CV validation.

**Figure 6.**
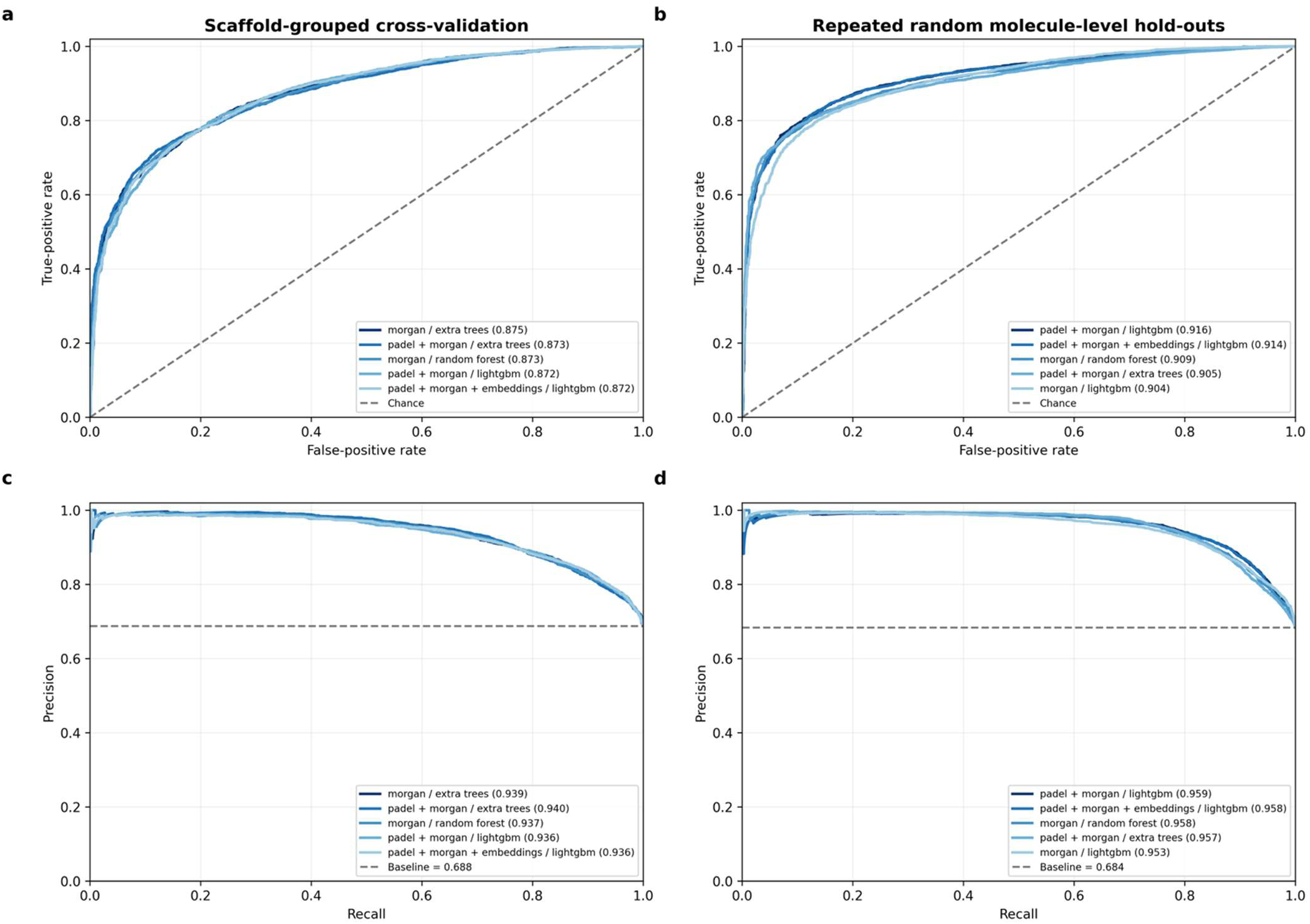
**(a) (b)** AUROC and AUPRC validation performance curves for best-performing models validated using scaffold CV, **(c) (d)** AUROC and AUPRC validation performance curves of best-performing models validated using repeated random molecule-level holdout validation.

### Controlled 3D and quantum-chemistry benchmark

We compared the PaDEL plus Morgan representation with two additional molecular representations using the same molecules, scaffold splits, external validation set, XGBoost classifier, and hyperparameter search procedure. The first configuration added 19 GFN2-xTB quantum-chemistry features to the PaDEL and Morgan features, while the second used a frozen 512-dimensional Uni-Mol v1 representation. This matched design allowed us to assess whether the additional quantum-chemistry or three-dimensional information improved prediction without differences in the evaluated molecules or validation procedure affecting the comparison. The PaDEL plus Morgan plus GFN2-xTB configuration produced the highest numerical performance, with an internal nested scaffold balanced accuracy of 0.780 and an external balanced accuracy of 0.670. However, this was only a small improvement over PaDEL plus Morgan. Internally, the addition of GFN2-xTB increased balanced accuracy by 0.001 and AUPRC by 0.002. Externally, balanced accuracy increased by 0.008 and AUPRC by 0.017. None of these differences remained statistically significant after Holm adjustment. The results therefore suggest that the GFN2-xTB features may provide some additional information, but they do not demonstrate a consistent improvement over the existing descriptor and fingerprint representation.

The frozen Uni-Mol representation did not improve performance. Compared with PaDEL plus Morgan, its internal balanced accuracy was lower by 0.041 and its internal AUPRC was lower by 0.036, with both differences remaining significant after Holm adjustment. External balanced accuracy was lower by 0.025 and external AUPRC was lower by 0.061, although these external differences were not statistically significant after adjustment. Complete results, scaffold-clustered 95% confidence intervals, raw and Holm-adjusted bootstrap p-values, McNemar tests, and prediction-level comparisons are provided in Supplementary Table S2 and the project repository.

Based on these results, the GFN2-xTB and Uni-Mol configurations were not added to the prediction tool. GFN2-xTB requires conformer generation and a quantum-chemistry calculation for every molecule, which would increase prediction time and computational requirements without a demonstrated improvement in performance. The frozen Uni-Mol representation also requires an additional software environment and greater computational resources, while performing less consistently than the PaDEL plus Morgan representation in this study. GFN2-xTB and Uni-Mol are presented as research benchmarks rather than replacement models. Future versions can reconsider their integration if BBB-specific fine-tuning, improved conformer treatment, or additional validation demonstrates a clear and consistent benefit.

Figure 5 summarizes balanced accuracy for the original model configurations. Figure 6 shows AUROC and precision-recall curves. The internal AUPRC baseline is approximately 0.688 because BBB+ is the majority class. The curves are interpreted together with balanced accuracy, sensitivity, specificity, MCC, scaffold separation, and external performance rather than as stand-alone evidence of generalization.

The PaDEL+Morgan LGBM and Extra Trees models and the PaDEL + Morgan + Embeddings XGBoost model were chosen to be incorporated in the prediction tool. For each configuration, the model was refitted on the full deduplicated training dataset using selected feature representations, preprocessing rules, and hyperparameters. Here, it is important to note that the top models produced numerically different metrics, but did not show statistically significant differences between them as tested by Wilcoxon paired testing. As no statistically significant differences were detected among the leading models, we selected the models to be included in the prediction tool based on a combination of numerical performance, robustness across validation techniques used, and practical deployment considerations. Models that were trained using only BBB-relevant descriptors performed worse than the models trained using all descriptors. These models are not used in the prediction tool currently, but they help to provide insight into how molecular descriptors can help to predict permeability. Performance metrics of these models and statistical comparison tables are included in the GitHub repository.

Source-exclusive validation trained on molecules found only in one source and tested on molecules found only in the other. Results were: PaDEL + Morgan, B3DB-exclusive to MoleculeNet BBBP-exclusive (training n = 5,941; test n = 280): balanced accuracy 0.754 and AUPRC 0.827; PaDEL + Morgan, MoleculeNet BBBP-exclusive to B3DB-exclusive (training n = 280; test n = 5,941): balanced accuracy 0.637 and AUPRC 0.778; PaDEL + Morgan + GFN2-xTB, B3DB-exclusive to MoleculeNet BBBP-exclusive (training n = 5,941; test n = 280): balanced accuracy 0.762 and AUPRC 0.833; PaDEL + Morgan + GFN2-xTB, MoleculeNet BBBP-exclusive to B3DB-exclusive (training n = 280; test n = 5,941): balanced accuracy 0.637 and AUPRC 0.783; Frozen Uni-Mol v1, B3DB-exclusive to MoleculeNet BBBP-exclusive (training n = 5,941; test n = 280): balanced accuracy 0.744 and AUPRC 0.831; Frozen Uni-Mol v1, MoleculeNet BBBP-exclusive to B3DB-exclusive (training n = 280; test n = 5,941): balanced accuracy 0.636 and AUPRC 0.793. The MoleculeNet-exclusive training direction is a small-cohort sensitivity analysis and is not treated as a stable deployment estimate. The reduction relative to internal scaffold validation supports the concern that source and endpoint heterogeneity remains important.

The top 10 most important features for the PaDEL+Morgan LGBM model and their highest-correlation pairs are shown in Table 4. This highlights the impact of removing highly correlated descriptors, particularly within PaDEL-derived features, which are known to contain significant redundancy. Eliminating these redundancies allows the models to focus on more informative and non-overlapping representations of molecular properties.

**Table 4.**
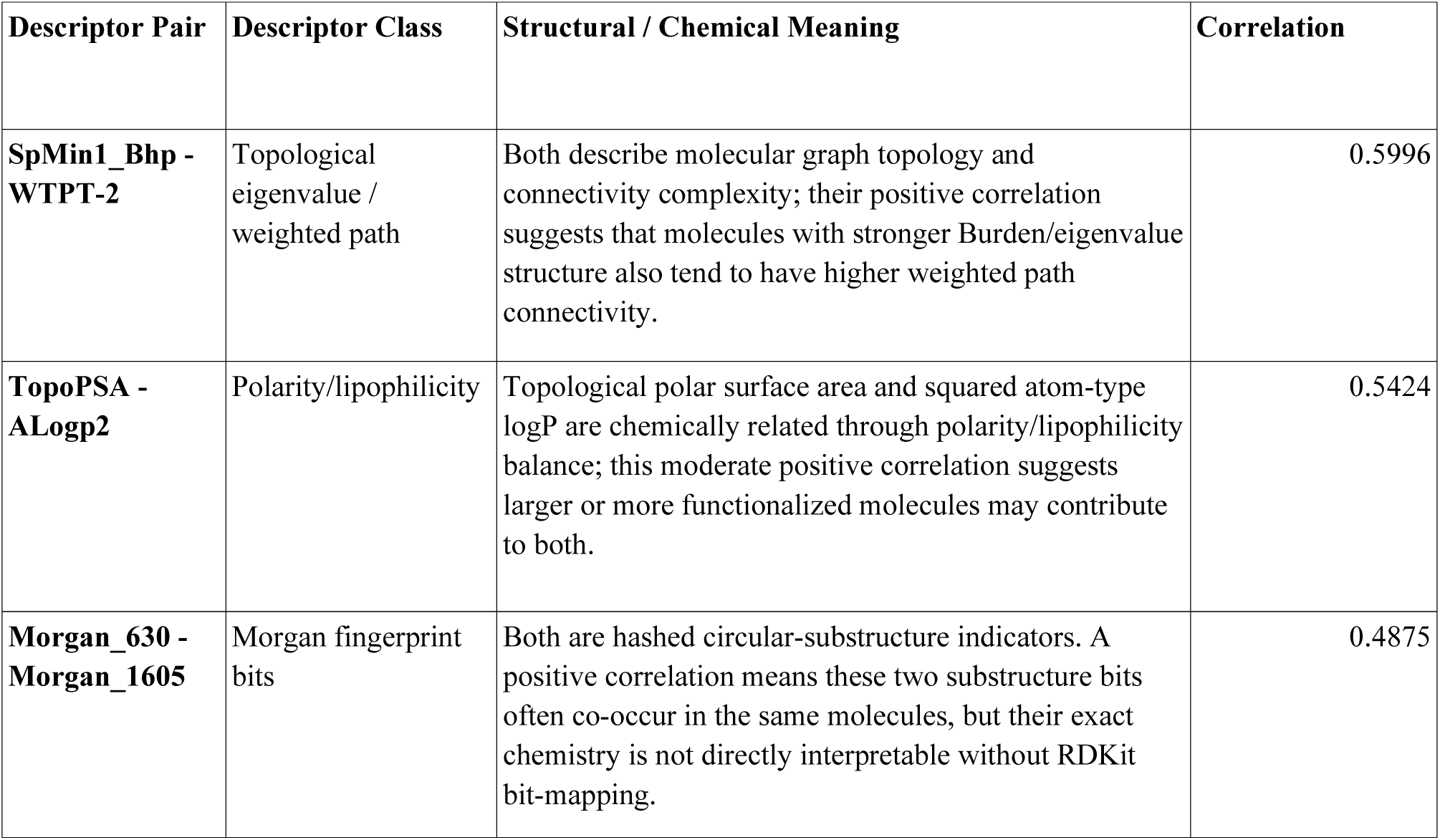
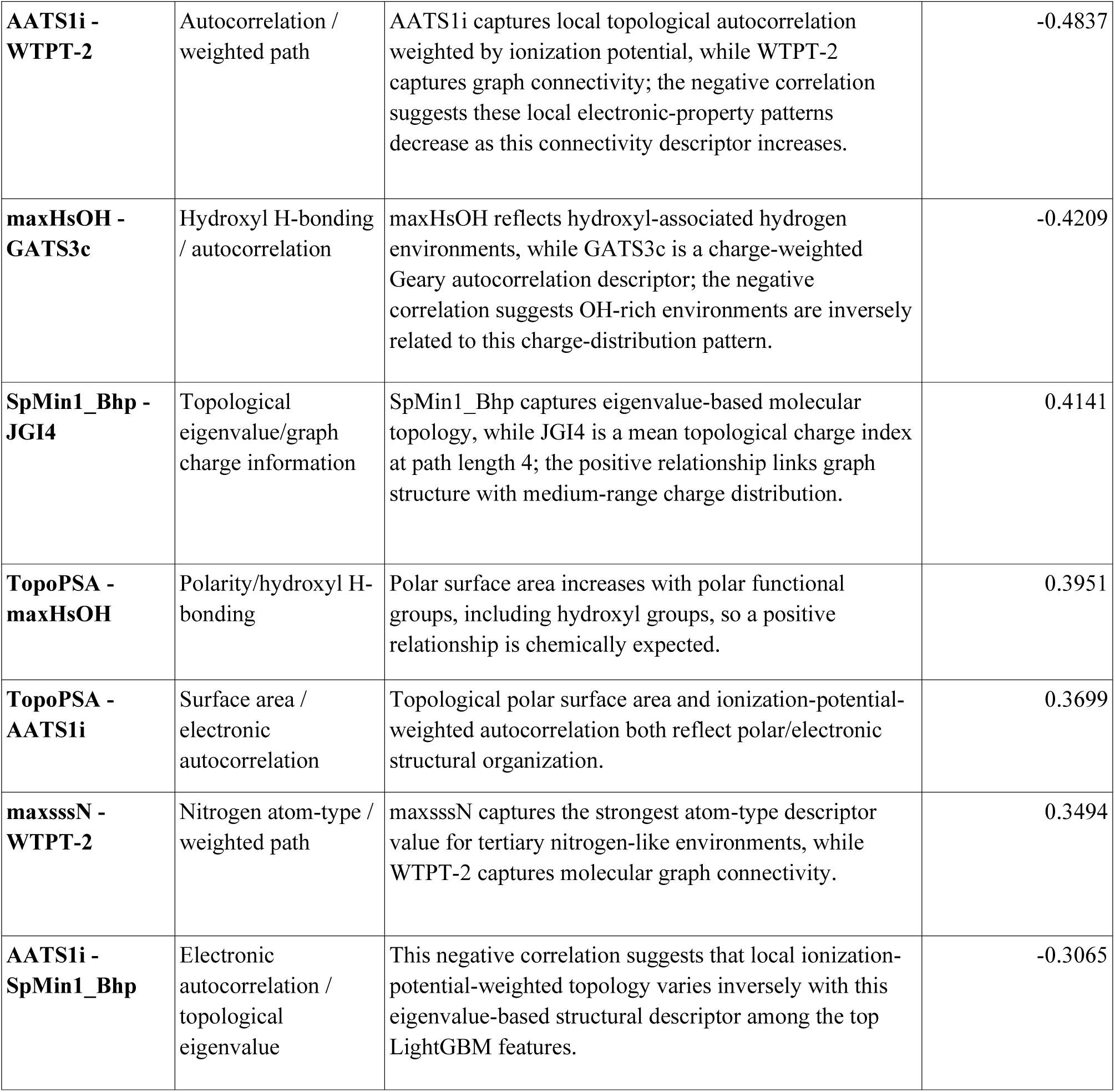
Descriptor correlation interpretation.

Figure 7(a) shows the SHAP values (Lundberg & Lee, 2017; Shapley, 1953) for the top features for the models chosen to be used in the BrainRoute prediction tool, helping to show which features contributed to push the model’s prediction toward the target class, which was set to be BBB permeable here. Figure 7(b) shows the correlation heatmap of the top 15 features for the PaDEL+Morgan+Embeddings XGBoost model. The strongest relationships were observed between TopoPSA and WTPT-3/WTPT-4, indicating that molecules with greater polar surface area also tended to have greater heteroatom and oxygen-based topological connectivity. The heatmap shows that models rely heavily on descriptors related to functional group distribution and intermolecular interaction potential. Feature importance information for all trained models is available in the provided GitHub repository. Table 5 shows the overview of the current literature in the field of BBB permeability prediction.

**Figure 7.**
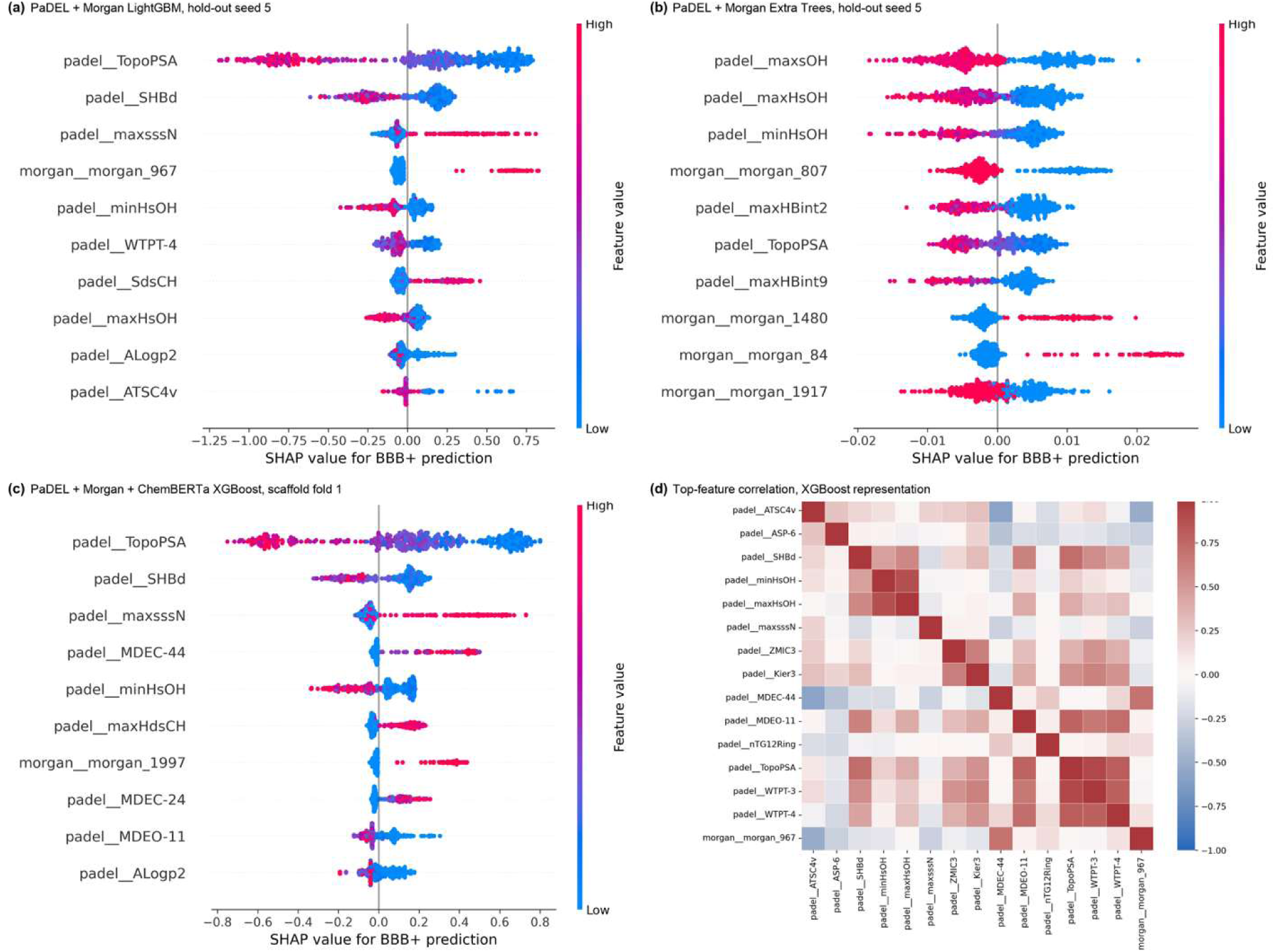
SHAP value distributions for the selected deployment models: **(a)** PaDEL + Morgan LightGBM, **(b)** PaDEL + Morgan Extra Trees, and **(c)** PaDEL + Morgan + ChemBERTa XGBoost. **(d)** Correlation heatmap of the top features for PaDEL + Morgan + ChemBERTa embeddings XGBoost model

**Table 5.** Contextual overview of BrainRoute with other studies exploring Blood-Brain Barrier permeability. Evaluations were not conducted on the same test set and are not directly comparable.

| Study | Data set | Model type/architecture | Reported metrics |
|---|---|---|---|
| Yuan et al. (2018) | 1,990 compounds; BBB+ = 1,550, BBB− = 440 | SVM with RBF kernel | Accuracy = 93.96%, sensitivity = 94.3%, specificity = 91.0%, MCC = 0.84 |
| Wang et al. (2018) | 2,358 compounds | SVM and kNN | Accuracy = 96.6%, sensitivity = 92.5%, specificity = 89.9% |
| Miao et al. (2019) | 462 compounds; BBB+ = 250, BBB− = 212 | Deep learning classifier | Accuracy = 97%, AUC = 0.98, F1 = 0.92 |
| Alsenan et al. (2020) | 2,342 compounds | RNN | Accuracy = 96.53%, sensitivity = 94.91%, specificity = 98.09%, MCC = 0.931, AUC = 0.986 |
| Shaker et al. (2020) | 7,162 compounds; BBB+ = 5,453, BBB− = 1,709 | LightGBM | Accuracy = 90%, sensitivity = 85%, specificity = 94% |
| Singh et al. (2020) | 605 training compounds + 1,566 validation compounds | RF, multilayer perceptron, SMO | Accuracy = 86.5% |
| Saxena et al. (2021) | 1,978 compounds; BBB+ = 1,550, BBB− = 440 | SVM with RBF kernel | Accuracy = 96.77%, AUC = 0.964, F1 = 0.975 |
| Kumar et al. (2022) | 3,605 total; BBB+ = 2,607, BBB− = 998; training = 2,704 | DNN | Accuracy = 98.07%, AUC = 0.992, AUPRC = 0.997, F1 = 0.987 |
| Mauri and Bertola (2022) | 3,884 curated; training = 3,525; BBB+ = 2,257, BBB− = 1,268 | KNN models + consensus model | Consensus test accuracy = 0.827, sensitivity = 0.760, specificity = 0.916 |
| Huang et al. (2024) | B3DB: 7,807 compounds; BBB+ = 4,956, BBB− = 2,851; CMUH: 2,499 compounds; BBB+ = 105, BBB− = 2,394 | MegaMolBART embeddings + XGBoost | Mixed dataset AUC = 0.88; B3DB-only AUC up to 0.90 |
| BrainRoute | 7,888 unique molecules<br>(Morgan: 7,888 x 2,048<br>ChemBERTa: 7,888 x 384<br>PaDEL-aligned: 7,882 x 1,874 | KNN + XGBoost + ET + RF + LGBM + LR | Primary scaffold CV BA 0.783; external ensemble BA 0.659, AUPRC 0.670 |

### Platform implementation and User Interface

The implemented platform workflow was tested using the following two molecules: D-Glucose and Dopamine Hydrochloride (Martel et al., 1996; Simpson et al., 1999). We chose these molecules for the implementation to capture a range of possibilities and how the workflow can be utilized even when dealing with molecules using active transport. These molecules were not present in the training data. Figure 8(a) shows the implementation workflow, where we started off with the BrainRoute prediction tool to get a prediction for the BBB permeability of these molecules. These predictions are automatically added to the database with the “predicted” tag. Then we simulated a user environment where another user can access the database platform and verify this information through the verification for submission and credibility architecture. The simulated user here provided a valid DOI to verify the prediction for both cases. In case 1 with Dopamine Hydrochloride, the provided user submission was used to verify that the model predictions were correct. In case 2, the user’s submission was to verify that the model prediction was incorrect. The changes are then reflected in the BrainRoute database platform, with these molecules now having a “verified” tag.

**Figure 8.**
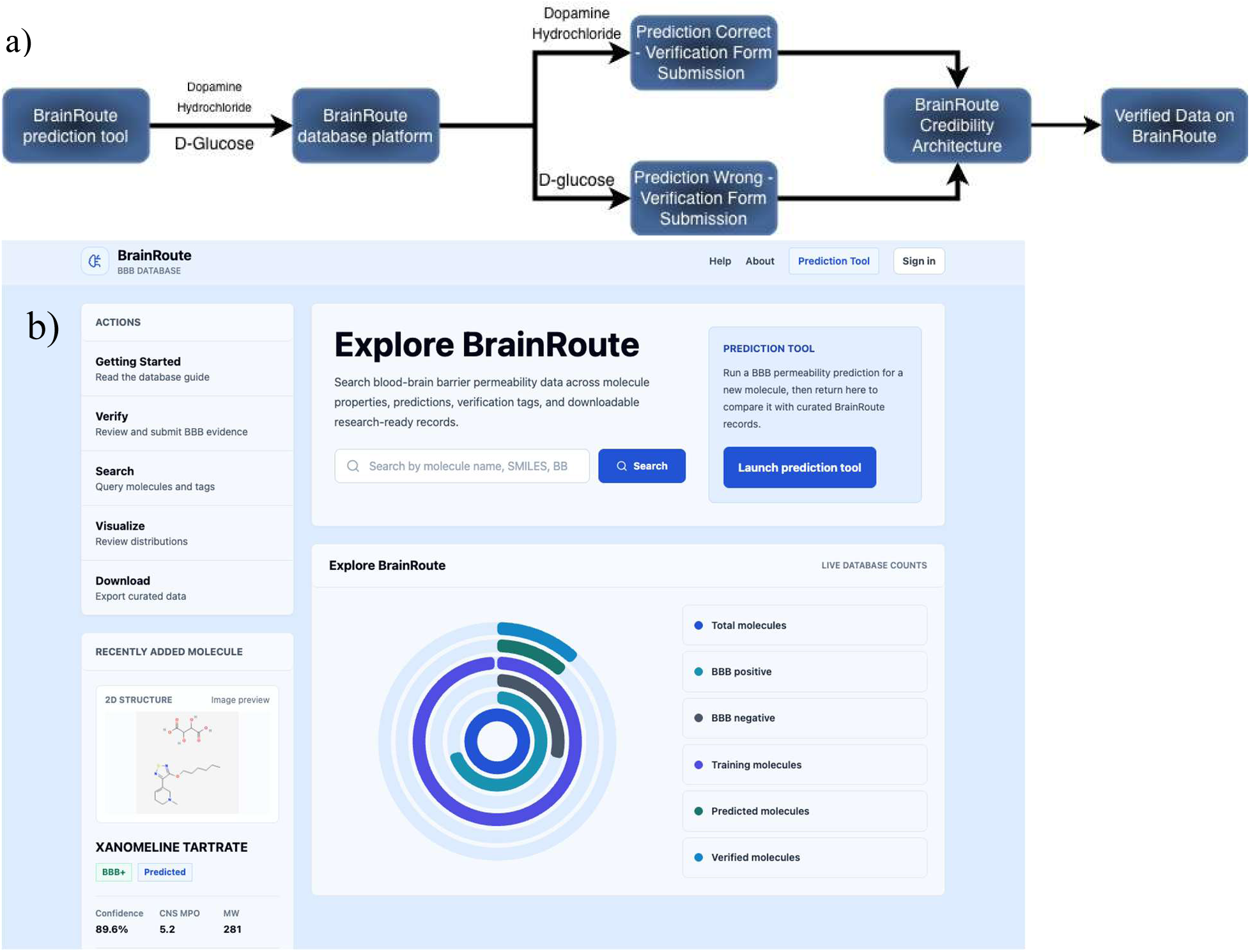
**(a)** Flowchart explaining platform verification protocol and information flow. **(b)** BrainRoute platform home page user interface.

External validation used the independent BBB dataset from Faramarzi et al. (2022). InChIKey overlap with the original training cohort was removed before evaluation. Of 921 retrieved external records, 358 exact training overlaps were removed and 153 additional records lacked complete PaDEL output, leaving 410 molecules, including 158 BBB+ and 252 BBB-. The overlap exclusions were 75.4% BBB+, whereas the PaDEL-failure exclusions were 32.7% BBB+ and were larger and more polar on average than retained molecules. The retained set had mean molecular weight 375.7 Da, mean logP 3.13, mean TPSA 80.36 square angstroms, mean nearest-training Tanimoto similarity 0.383, median similarity 0.315, and 84 molecules with a non-empty Murcko scaffold observed in training. These differences show that attrition was not random and limit the scope of the external estimate.

On the 410-molecule external cohort, the deployed ensemble achieved balanced accuracy 0.659 (95% CI 0.600-0.712) and AUPRC 0.670 (95% CI 0.583-0.752). Individual model metrics and scaffold-clustered confidence intervals are reported in Supplementary Table S3. All external validation predictions, datasets, and performance summaries are provided in the GitHub repository.

### Case Study

To best illustrate how BrainRoute can help with CNS drug discovery and research, we have chosen inhibitors of the mechanistic Targets of Rapamycin (mTOR), colloquially known by researchers as mTOR inhibitors, as the primary case study because of their important role in modulating cellular cleaning mechanisms, which can help with neurodegenerative diseases. The mechanistic Target of Rapamycin (mTOR) is a serine-threonine protein kinase that operates as a regulator of a vast array of pivotal cellular mechanisms, such as cellular metabolism, integrating signals from nutrients, growth factors, and energy status to control critical processes, including cell growth, proliferation, survival, protein synthesis, and lysosomal biogenesis. (Davoody *et al.,* 2023; Querfurth and Lee, 2021; Ather *et al.,* 2018). Most notably, it is a key player in autophagy - a cellular method via the formation of a structure known as an autophagosome around the protein and organelles, which are taken to be disposed of by the lysosomes (Heras-Sandoval, Pérez-Rojas, and Pedraza-Chaverri, 2020). This is a particularly important facet of the mTOR pathway, as both the hypo- and hyperactivation of autophagy are associated with neurodegeneration, so its functional regulation is crucial. (Dolcetta and Dominici, 2019; Heras-Sandoval, Pérez-Rojas and Pedraza-Chaverri, 2020; Aribi *et al.,* 2025).

The process of autophagy can be considered a cellular “self-cleaning” mechanism. As neurons are highly dependent on efficient protein quality control, being long-lived post-mitotic cells, dysfunction of the autophagy process can lead to accumulation of misfolded protein aggregates, which are linked to neurodegenerative conditions such as Alzheimer’s disease (AD).

Hyperactivation of mTORC1 (mTOR Complex 1, a kinase complex that regulates growth, metabolism, and autophagy) is correlated with suppressed autophagy, leading to dysfunctional clearance of amyloid-*β* (A*β*) and hyperphosphorylated tau aggregates, which are key pathological features of AD. Furthermore, dysregulated mTORC1 signaling is also linked to cerebrovascular impairments, including BBB breakdown and endothelial integrity compromise in preclinical models of AD. Hyperactivation of mTOR has also been strongly implicated in a myriad of pathologies associated with aging (Davoody *et al.,* 2023). In the nervous system, many chronic diseases and pathological aging processes are driven by oxidative stress, inflammation, mitochondrial failure, and insulin resistance, all of which intensify due to mTOR hyperactivation. (Bouyahya *et al.,* 2022; Querfurth and Lee, 2021).

This makes mTOR a compelling therapeutic target, and since the discovery of the prototypical inhibitor, rapamycin, it has only recently moved from its initial use as an immunosuppressant to a target in the intersecting field of neuroscience and anti-aging. (Lamming *et al.,* 2013; Davoody *et al.,* 2023; Ben Aribi *et al.,* 2024; Ben Aribi *et al.,* 2025; El Abed *et al.,* 2023; Alaya *et al.,* 2024). Experimental and preclinical neurodegeneration models have shown that rapamycin and its analogs clear the protein aggregates, restore cerebral blood flow, and preserve BBB integrity (Van Skike *et al.,* 2018; Uddin *et al.,* 2020). Furthermore, many AD mouse models have shown that mTORC1 inhibition has increased autophagy (Figure 9), reduced A*β* and tau pathology, and improved synaptic plasticity and cognition (Cordos *et al.,* 2025).

**Figure 9.**
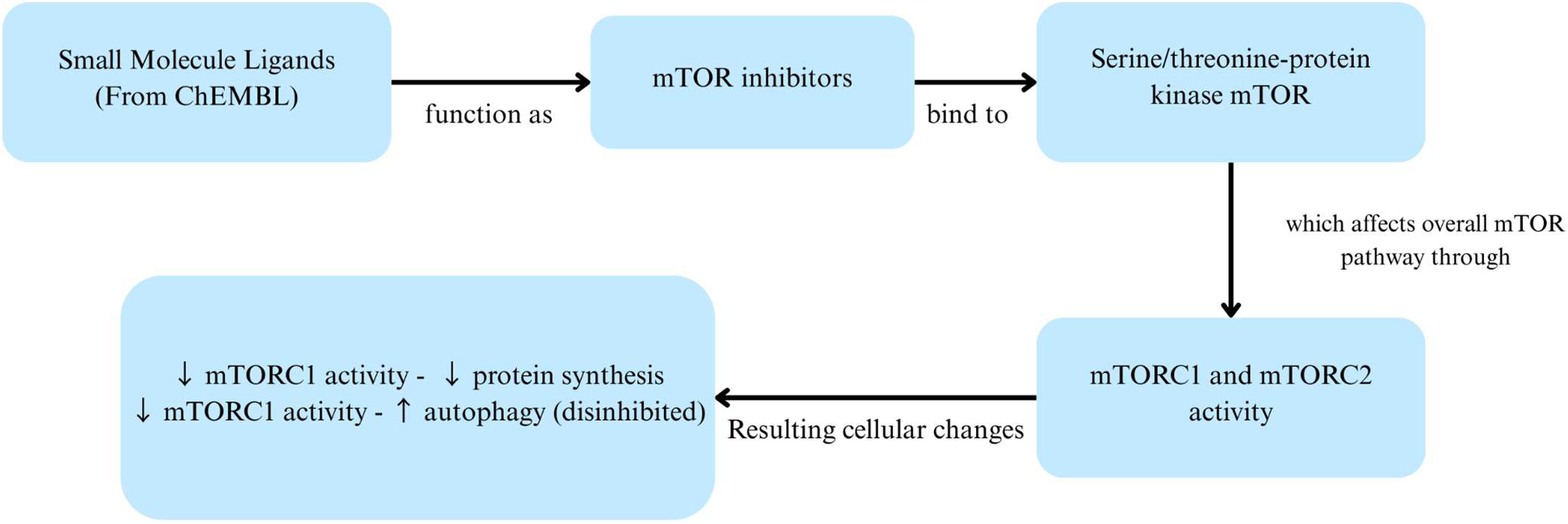
Conceptual flowchart illustrating the role of mTOR inhibitors in autophagy regulation. Small molecules analyzed in this study can bind to serine/threonine-protein kinase mTOR, inhibiting the kinase’s activity. This can result in a reduction of mTORC1 signaling, which can increase autophagic activity and enhance the clearance of misfolded proteins. These inhibitors can be potentially relevant for neurodegenerative disease drug discovery applications. (Rubinsztein *et al.,* 2011; Querfurth & Lee, 2021; Saxton & Sabatini, 2017).

However, rapamycin’s translation to clinical practice is hindered by its myriad unpleasant side effects - from triggering system-wide inflammation, nephrotoxicity, and blunting the body’s natural healing process via immunosuppression (Querfurth and Lee, 2021). As such, the discovery and development of safer rapalogs has emerged as a growing area of research, with active compounds ranging from peptides, antibiotics, and analgesics to natural extracts, all of which have promising mTOR-modulating potential (Fakhri *et al.,* 2021; Heras-Sandoval, Pérez-Rojas, and Pedraza-Chaverri, 2020). By integrating insights from novel drug discovery and ancient traditional medicinal herbs, a potentially safe rapalog may be achievable in the near future (Pouria Mosaddeghi *et al.,* 2021; Heras-Sandoval, Pérez-Rojas, and Pedraza-Chaverri, 2020; Bouyahya *et al.,* 2022). These rapalogs may not only change the course of disease management for neurodegenerative diseases but also for ocular degenerative pathologies such as glaucoma, retinal pigmentosa, diabetic retinopathy, and age-related macular degeneration, autoimmune diseases, and oncological care (Wang *et al.,* 2022; Mueed *et al.,* 2019).

The case-study contains 26 ChEMBL mechanism records annotated with target CHEMBL2842, target type SINGLE PROTEIN, organism Homo sapiens, and action type INHIBITOR. The records span maximum development phases 1 to 4. No assay-level filter or potency threshold was applied because these were mechanism records rather than an activity query. Twenty-two records had non-missing SMILES and were processed by the BrainRoute batch interface. They are described as clinical-development mTOR inhibitor records, not as 22 approved drugs.

**Table 6.** Serine/threonine-protein kinase mTOR inhibitors predictions using BrainRoute.

| ChEMBL ID | Prediction (BBB+/-) | Confidence | Uncertainty | Agreement |
|---|---|---|---|---|
| CHEMBL1258517 | BBB- | 58.99 | 9.21 | 2/3 (66.7%) |
| CHEMBL5095025 | BBB- | 71.9 | 16.61 | 3/3 (100.0%) |
| CHEMBL2326966 | BBB- | 90.01 | 10.69 | 3/3 (100.0%) |
| CHEMBL1236962 | BBB- | 85.62 | 12.35 | 3/3 (100.0%) |
| CHEMBL2141712 | BBB- | 66.14 | 9.64 | 3/3 (100.0%) |
| CHEMBL2331680 | BBB+ | 59.25 | 3.21 | 2/3 (66.7%) |
| CHEMBL592445 | BBB- | 79.29 | 12.66 | 2/3 (66.7%) |
| CHEMBL1234354 | BBB- | 83.12 | 15.53 | 3/3 (100.0%) |
| CHEMBL1922094 | BBB- | 71.37 | 12.79 | 3/3 (100.0%) |
| CHEMBL3586573 | BBB- | 64.46 | 11.39 | 3/3 (100.0%) |
| CHEMBL75880 | BBB- | 86.85 | 10.05 | 3/3 (100.0%) |
| CHEMBL3545097 | BBB- | 69.75 | 8.72 | 2/3 (66.7%) |
| CHEMBL2103839 | BBB- | 65.85 | 9.13 | 3/3 (100.0%) |
| CHEMBL1801204 | BBB- | 52.48 | 12.72 | 2/3 (66.7%) |
| CHEMBL3586404 | BBB- | 59.85 | 1.98 | 2/3 (66.7%) |
| CHEMBL3120215 | BBB- | 74.26 | 4.65 | 3/3 (100.0%) |
| CHEMBL3393066 | BBB+ | 67.29 | 8.09 | 2/3 (66.7%) |
| CHEMBL2336325 | BBB- | 57.97 | 4.65 | 2/3 (66.7%) |
| CHEMBL1879463 | BBB- | 68.95 | 10.42 | 2/3 (66.7%) |
| CHEMBL3218578 | BBB- | 69.33 | 7.63 | 3/3 (100.0%) |
| CHEMBL3545366 | BBB- | 66.7 | 10.58 | 2/3 (66.7%) |
| CHEMBL4297181 | BBB- | 50.94 | 14.12 | 2/3 (66.7%) |

The platform screening classified 2 of the 22 mTOR inhibitor records as BBB+ and 20 as BBB- (Table 6). These are computational outputs from the Version 1 models, not experimental permeability measurements. Given the deployed ensemble external balanced accuracy of approximately 0.66, the case study demonstrates the platform workflow and generates hypotheses for experimental testing; it does not support a conclusion that only two compounds cross the BBB.

## 4. Discussion

The clearest evidence of how BrainRoute’s prediction tool is likely to perform in practice comes from the independent external validation set rather than from internal cross-validation alone. On 410 molecules with no InChIKey or scaffold overlap with the training cohort, the deployed three-model ensemble achieved a balanced accuracy of 0.659 (95% CI 0.600–0.712) and an AUPRC of 0.670 (95% CI 0.583–0.752). We treat this external estimate as the primary evidence of the tool’s generalization, because chemical similarity between training and test molecules can substantially inflate internal validation metrics. Consistent with this concern, internal balanced accuracy varied considerably by validation strategy: the highest mean balanced accuracy under repeated random 80/20 molecule-level hold-outs was 0.836 ± 0.008 (PaDEL + Morgan LightGBM), whereas the primary five-fold Bemis-Murcko scaffold-grouped cross-validation, which assigns molecules sharing a core scaffold entirely to one partition, produced a lower mean balanced accuracy of 0.783 ± 0.026 (PaDEL + Morgan + ChemBERTa XGBoost). The gap between these two internal estimates, and the further reduction observed on external validation, together indicate that performance figures drawn from conventional random splits should not be read as estimates of real-world generalization for BBB permeability models. We therefore treat the scaffold-grouped result as the appropriate internal benchmark and the external result as the figure that should guide expectations about deployment performance.

Dataset heterogeneity, assay differences, and source-specific label definitions are plausible contributors to this external drop, consistent with the non-random attrition and reduced chemical similarity observed in the external cohort. Source-exclusive validation, in which models were trained on molecules from one source and tested only on molecules exclusive to the other, showed a comparable reduction relative to internal scaffold validation, reinforcing that source and endpoint heterogeneity remains an important open challenge for BBB permeability modeling generally, not only for this dataset.

These outputs should be interpreted as computational screening results rather than definitive evidence of BBB transport. Descriptor-based models of this kind cannot fully capture active transport, transporter-mediated uptake or efflux, or other biological mechanisms that influence BBB permeability (Cornelissen et al., 2023; Kusuhara et al., 1997; Löscher & Potschka, 2005), a limitation illustrated directly by the case-study molecule D-glucose, which crosses the BBB through a dedicated transporter rather than passive diffusion. The controlled comparison against GFN2-xTB quantum-chemistry descriptors and a frozen Uni-Mol v1 representation did not identify a representation that consistently outperformed the deployed PaDEL plus Morgan features under matched molecules, splits, and evaluation procedure; this is discussed further as a limitation and future direction below.

To give more context about the models used for the prediction tool, we present a comparative analysis with commonly used metrics reported from previously reported BBB prediction platforms (Table 5). However, it is important to note here that these comparisons are not conducted on a shared, externally held-out test set. Many existing platforms do not provide a clear user interface, reproducible pipelines, publicly available models, or sufficient implementation details, leading to limitations with independent evaluations under standardized conditions. Hence, the values reported were derived from supporting published articles for previous work and should be interpreted as qualitative rather than strictly quantitative comparisons. This also highlights the challenge of reproducibility in the field and shows the importance of benchmarking frameworks for future research (Pineau et al., 2021).

BrainRoute provides a community-driven platform for BBB research that integrates interactive data exploration, verification workflows, and continuously expanding molecular information. The established authentication and validation framework provides a seamless interaction experience to collect and contribute data related to BBB permeability. Through the implementation example, we can see how the platform can provide utility not just in predicting blood-brain barrier permeability but also in providing an interface to verify and curate research findings. This design allows BrainRoute to work as an ever-changing and guiding resource that can keep up with and contribute to the latest research methodologies by providing data and verified insights.

### 4.1 Limitations

Through this study, we successfully developed an integrated BBB prediction and database platform to support future BBB research. However, several limitations should be considered when interpreting its utility. The prediction tool relies mainly on conventional machine-learning classifiers trained using molecular descriptors and fingerprints. These models capture physicochemical and structural trends associated with passive diffusion but do not directly represent active transport, transporter-mediated uptake or efflux, membrane-protein interactions, dose, formulation, species differences, or disease state. Compounds such as D-glucose, which cross the BBB through specific transporter mechanisms, may therefore not be represented accurately using molecular properties alone. This reflects a broader limitation of descriptor-based prediction, as complex biological transport cannot be fully inferred from molecular structure (Pardridge, 2012). Computational predictions should consequently be interpreted as screening-level evidence rather than a replacement for in vitro or in vivo assessment.

The controlled benchmark extended the analysis to GFN2-xTB quantum-chemistry descriptors and frozen Uni-Mol v1 representations, but these approaches also have limitations. Uni-Mol was used as a frozen 512-dimensional representation rather than being fine-tuned specifically for BBB permeability because the available system had 16 GB of unified memory and no supported dedicated GPU. The GFN2-xTB descriptors were calculated from one force-field-minimized conformer using a gas-phase single-point calculation. Conformer ensembles, solvent effects, pH-dependent microspecies, and higher-level quantum calculations were outside the available computational scope. Molecules that did not produce a successful xTB calculation or a valid three-dimensional Uni-Mol conformer were excluded from all three matched benchmark configurations. This maintained a fair comparison but reduced the range of molecules represented. Neither approach demonstrated sufficient improvement to be used with production models, although BBB-specific fine-tuning and more complete conformational and quantum-chemical treatment can be evaluated in future versions.

The dataset also contains limitations associated with combining BBB labels from different sources and evidence types. The download dates were not retained during dataset assembly. The exact files used in this study are therefore documented through their repository copies, record counts, SHA-256 checksums, and row-level source mappings. These records identify the data used for model development but cannot recover source information that was not retained during the initial assembly.

Finally, the community verification workflow currently depends on manual review by the development team. Manual curation allows submissions and supporting evidence to be evaluated in context, but it introduces human dependency and will become difficult to maintain as the platform grows. Future versions can add automated checks for molecular identity, required metadata, source consistency, duplicate records, and conflicting labels while retaining human review for evidence interpretation and final verification. BrainRoute is therefore positioned as a complementary and developing resource that emphasizes integration, accessibility, and transparent data review rather than as a replacement for experimental BBB evaluation.

### 4.2 Future work

Future development of BrainRoute follows two complementary tracks: continued development of the platform itself, and a dedicated effort to improve the underlying predictive models beyond the scope of the present release.

On the modeling side, the present study represents a first, platform-oriented iteration rather than a state-of-the-art predictive benchmark, and several directions follow directly from its limitations. The frozen Uni-Mol v1 representation and the single-conformer GFN2-xTB descriptors evaluated in this study were constrained by the available CPU-only hardware; we plan to revisit both approaches under GPU resources that support end-to-end Uni-Mol fine-tuning, multi-conformer ensembles, and higher-level quantum-chemical treatment, since neither representation has yet been given a fair opportunity to demonstrate its potential benefit. We also considered more recent quantum-chemistry-aware learned representations, such as qcGEM (Wang & Gong, 2025) as alternatives to a classical descriptor-based quantum-chemistry featurization. These approaches require either large-scale pretraining corpora or GPU-based encoding architectures that were not compatible with the CPU-only hardware available for this study, so we instead used the lighter-weight, fully reproducible GFN2-xTB descriptor set as a first, tractable step toward incorporating quantum-chemical information. We also plan to evaluate the deployed ensemble under class distributions weighted toward BBB-molecules, more representative of prospective screening libraries than the BBB+-enriched curated datasets used for training and evaluation here. Because these modeling improvements require a depth of methodological development, benchmarking, and validation that would be difficult to accommodate within a platform-focused manuscript, we intend to report on them separately in a dedicated, model-focused publication, allowing this modeling work to be evaluated on its own terms while the present study establishes the data infrastructure that future models can be deployed into.

On the platform side, we plan to incorporate a fine-tuned LLM tailored for CNS drug discovery and research in the platform. This would help users to understand more about individual molecular properties through integration with the current molecule card design, as well as use it for faster filtering. We also plan to fine-tune the ChemBERTa-77M-MLM model on BBB permeability data to better utilize the model’s capabilities. Additionally, we aim to introduce automated checks for molecular identity, required metadata, source consistency, duplicate records, and conflicting labels within the verification workflow, reducing dependence on fully manual review as submission volume grows, while retaining human review for evidence interpretation and final verification.

Furthermore, we aim to include more filtering options, scoring metrics, and better integration with existing external datasets and published resources to expand the platform’s potential analytical use cases. We plan to make BrainRoute a comprehensive, centralized resource for all BBB-related data.

## 5. Conclusion

This work introduces BrainRoute, a platform representing a significant advancement in predicting blood–brain barrier permeability by unifying robust machine learning methodologies with data aggregation and community-driven validation within a single platform. Previous works with BBB-related datasets and predictive models lack standardization and provide limited mechanisms for validation and growth. BrainRoute addresses this gap by providing an integrated environment for molecular prediction and contributing experimental evidence. This design positions the platform as both a research tool and a collaborative knowledge infrastructure for the broader scientific community.

The current implementation of the platform represents Version 1, with a clearly defined pathway for iterative improvement through expanded datasets, enhanced filtering, and incorporation of Artificial Intelligence (AI) tools. The platform aims to grow through community engagement and foster new model development and research with more data and reliable insights into BBB permeability.

## Declarations

### Ethics approval and consent to participate

Not applicable.

### Consent to Participate

Not applicable.

### Consent for publication

Not applicable.

### Clinical Trial Number

Not applicable.

### Availability of data and materials

The datasets analyzed during the current study are available in the BrainRoute GitHub repository, https://github.com/omicscodeathon/brainroute. The BrainRoute platform source code is available at https://github.com/omicscodeathon/brainroutedb. The versions presented in this manuscript correspond to commits 7cefeb054e268d7cfe81f2738d7bbc5715ac9d18 and 17da493e8173600bf72c1c9806676310bf3d548a, respectively. The data supporting the results reported in this manuscript are included within the article and the corresponding GitHub repository.

Prediction tool’s GitHub repository: https://github.com/omicscodeathon/brainroute Any restrictions on use by non-academics: None.

### Competing interests

The authors declare that they have no competing interests.

## Funding

The authors declare that no financial support was received for the research, authorship, or publication of this article.

## Authors’ contributions

SAS conceived the original idea. SAS developed the project pipeline, performed the computational analysis of the case-study data, and drafted the manuscript. SAS, LTB, LWC, and NEA contributed to writing, reviewing, and editing the manuscript. All authors contributed to the original draft. OIA administered and supervised the computational analysis, provided the resources required to complete the analysis, and provided guidance. OIA also edited and reviewed the final manuscript and provided critical feedback. All authors read and approved the final manuscript.

## Acknowledgements

The authors thank the National Institutes of Health Office of Data Science Strategy for its support before and during the October 2025 Omics Codeathon organized by the Institute for Genomic Medicine Research in collaboration with the African Society for Bioinformatics and Computational Biology.

## Generative AI disclosure

Generative AI tools were used to assist with code development for the machine learning models generated platform implementation. Authors have reviewed and verified all generated code, text, analyses, and results and take complete responsibility for the final work.

## List of Abbreviations

CNS: Central Nervous System
KNN: K-Nearest Neighbors
XGBoost: eXtreme Gradient Boosting
ET: Extra Trees Classifier
RF: Random Forest
LGBM: Light Gradient Boosting Machine
LR: Logistic Regression

## Notes

### Competing Interest Statement

The authors have declared no competing interest.

