## Supplementary Tables for "BrainRoute integrates machine learning prediction with community-driven verification of blood-brain barrier permeability"

### BrainRoute Supplementary Tables

#### Supplementary Table S1

Source endpoints, evidence fields, class distributions, and limitations.

| Source | Endpoint and mapping | Raw records | BBB+ | BBB- | Evidence fields and limitations |
|---|---|---|---|---|---|
| B3DB | BBB+/BBB- classification: The published B3DB categorical classification was retained; BBB+ was mapped to 1 and BBB- to 0 | 7,807 | 4,956 | 2,851 | source reference codes; evidence-quality group for all rows; numerical logBB for 1058 rows; source-specific threshold for 3621 rows. B3DB combines numerical, threshold-based, and categorical literature evidence; its quality groups must be considered when interpreting the binary endpoint |
| MoleculeNet BBBP | p_np: p_np=1 was mapped to BBB+ and p_np=0 was mapped to BBB- | 2,050 | 1,567 | 483 | compound name; SMILES; binary endpoint derived from the Martins et al. BBB penetration collection. The local file does not retain assay type, species, experimental method, or a numerical permeability value at row level |

Source reconstruction retained 9,844 RDKit-valid records representing 8,090 unique InChIKeys. Forty-eight InChIKeys had more than one source label. The modeling-table curation excluded 32 conflicting-label structures; the row-level audit identified 24 retained modeling molecules with disagreeing source records.

#### Supplementary Table S2

Matched nested-scaffold representation check. Values are estimates with scaffold-clustered 95% confidence intervals.

| Configuration | Internal balanced accuracy | Internal AUPRC | External balanced accuracy | External AUPRC |
|---|---|---|---|---|
| PaDEL + Morgan | 0.779 (0.751-0.803) | 0.932 (0.911-0.951) | 0.662 (0.605-0.714) | 0.672 (0.569-0.763) |
| PaDEL + Morgan + GFN2-xTB | 0.780 (0.756-0.803) | 0.934 (0.915-0.953) | 0.670 (0.613-0.723) | 0.690 (0.581-0.775) |
| Frozen Uni-Mol v1 | 0.737 (0.714-0.761) | 0.897 (0.870-0.922) | 0.637 (0.589-0.688) | 0.611 (0.512-0.702) |

Paired comparisons

| Cohort | Comparison | Metric | Difference | 95% CI | Holm-adjusted bootstrap p | Holm-adjusted McNemar p |
|---|---|---|---|---|---|---|
| Internal | PaDEL + Morgan + GFN2-xTB vs PaDEL + Morgan | Balanced accuracy | +0.001 | -0.007 to +0.009 | 1.0000 | 0.8390 |
| Internal | PaDEL + Morgan + GFN2-xTB vs PaDEL + Morgan | AUPRC | +0.002 | -0.001 to +0.005 | 0.6160 | 0.8390 |
| Internal | Frozen Uni-Mol v1 vs PaDEL + Morgan | Balanced accuracy | -0.041 | -0.060 to -0.022 | <0.001 | 0.0000 |
| Internal | Frozen Uni-Mol v1 vs PaDEL + Morgan | AUPRC | -0.036 | -0.045 to -0.027 | <0.001 | 0.0000 |
| External | PaDEL + Morgan + GFN2-xTB vs PaDEL + Morgan | Balanced accuracy | +0.008 | -0.012 to +0.028 | 1.0000 | 1.0000 |
| External | PaDEL + Morgan + GFN2-xTB vs PaDEL + Morgan | AUPRC | +0.017 | -0.002 to +0.039 | 0.5050 | 1.0000 |
| External | Frozen Uni-Mol v1 vs PaDEL + Morgan | Balanced accuracy | -0.025 | -0.079 to +0.029 | 1.0000 | 0.8390 |
| External | Frozen Uni-Mol v1 vs PaDEL + Morgan | AUPRC | -0.061 | -0.129 to +0.004 | 0.3600 | 0.8390 |

#### Supplementary Table S3

Independent external results for each deployed model and their probability-averaged ensemble. Confidence intervals use 2,000 scaffold-clustered bootstrap resamples.

| Model | Balanced accuracy | AUPRC | Sensitivity | Specificity | MCC | Brier score |
|---|---|---|---|---|---|---|
| PaDEL + Morgan LightGBM | 0.651 (0.594-0.699) | 0.651 (0.564-0.732) | 0.652 (0.572-0.725) | 0.651 (0.578-0.714) | 0.295 (0.184-0.393) | 0.222 (0.195-0.252) |
| PaDEL + Morgan Extra Trees | 0.652 (0.595-0.700) | 0.679 (0.590-0.755) | 0.665 (0.584-0.739) | 0.639 (0.560-0.711) | 0.296 (0.183-0.393) | 0.212 (0.197-0.229) |
| PaDEL + Morgan + ChemBERTa XGBoost | 0.655 (0.599-0.706) | 0.644 (0.549-0.735) | 0.620 (0.540-0.696) | 0.690 (0.618-0.753) | 0.306 (0.192-0.408) | 0.211 (0.189-0.237) |
| BrainRoute probability-averaged ensemble | 0.659 (0.600-0.712) | 0.670 (0.583-0.752) | 0.652 (0.568-0.726) | 0.667 (0.590-0.735) | 0.311 (0.194-0.417) | 0.209 (0.188-0.233) |

#### Supplementary Table S4

External-set attrition and chemical-space comparison.

| Cohort | n | BBB+ | BBB- | BBB+ % | Mean MW | Mean logP | Mean TPSA | Mean nearest-training Tanimoto | Seen non-empty scaffold |
|---|---|---|---|---|---|---|---|---|---|
| Exact training overlaps removed | 358 | 270 | 88 | 75.4 | 320.7 | 2.76 | 60.62 | Not calculated | 328 |
| PaDEL descriptor failures excluded | 153 | 50 | 103 | 32.7 | 532.5 | 3.24 | 118.59 | 0.427 | 36 |
| Retained external validation cohort | 410 | 158 | 252 | 38.5 | 375.7 | 3.13 | 80.36 | 0.383 | 84 |

#### Supplementary Table S5

Source-exclusive validation. The fixed classification threshold was 0.5.

| Configuration | Direction | Train n | Test n | Test BBB+ / BBB- | Balanced accuracy | AUPRC | Sensitivity | Specificity |
|---|---|---|---|---|---|---|---|---|
| PaDEL + Morgan | B3DB-exclusive → MoleculeNet-exclusive | 5,941 | 280 | 150 / 130 | 0.754 | 0.827 | 0.693 | 0.815 |
| PaDEL + Morgan | MoleculeNet-exclusive → B3DB-exclusive | 280 | 5,941 | 3909 / 2032 | 0.637 | 0.778 | 0.726 | 0.547 |
| PaDEL + Morgan + GFN2-xTB | B3DB-exclusive → MoleculeNet-exclusive | 5,941 | 280 | 150 / 130 | 0.762 | 0.833 | 0.740 | 0.785 |
| PaDEL + Morgan + GFN2-xTB | MoleculeNet-exclusive → B3DB-exclusive | 280 | 5,941 | 3909 / 2032 | 0.637 | 0.783 | 0.728 | 0.545 |
| Frozen Uni-Mol v1 | B3DB-exclusive → MoleculeNet-exclusive | 5,941 | 280 | 150 / 130 | 0.744 | 0.831 | 0.673 | 0.815 |
| Frozen Uni-Mol v1 | MoleculeNet-exclusive → B3DB-exclusive | 280 | 5,941 | 3909 / 2032 | 0.636 | 0.793 | 0.678 | 0.593 |

#### Supplementary Table S6

Model-specific handling of class imbalance and validation roles.

| Classifier | Imbalance handling |
|---|---|
| Logistic Regression | class_weight='balanced' |
| Random Forest | class_weight='balanced' |
| Extra Trees | class_weight='balanced' |
| LightGBM | class_weight='balanced' |
| XGBoost | scale_pos_weight = n_negative / n_positive in each outer training partition |
| K-Nearest Neighbors | no class-weight parameter; distance weighting was among tuned options |

| Validation analysis | Role | Splits | InChIKey overlap | Maximum scaffold overlap | Interpretation |
|---|---|---|---|---|---|
| Primary five-fold scaffold-grouped cross-validation | primary | 5 | 0 | 0 | Primary internal estimate; core scaffolds are separated |
| Secondary repeated random holdout on the InChIKey-deduplicated cohort | secondary | 5 | 0 | 579 | Secondary analysis; structurally related molecules may occur across partitions |
| Secondary scaffold-group holdout | secondary | 1 | 0 | 0 | Secondary analysis; structurally related molecules may occur across partitions |

Supporting files are located in BrainRoute model repository.
